# Light-regulated collective contractility in a multicellular choanoflagellate

**DOI:** 10.1101/661009

**Authors:** Thibaut Brunet, Ben T. Larson, Tess A. Linden, Mark J. A. Vermeij, Kent McDonald, Nicole King

## Abstract

Collective cell contractions that generate global tissue deformations are a signature feature of animal movement and morphogenesis. Nonetheless, the ancestry of collective contractility in animals remains mysterious. While surveying the Caribbean island of Curaçao for choanoflagellates, the closest living relatives of animals, we isolated a previously undescribed species (here named *Choanoeca flexa* sp. nov.), that forms multicellular cup-shaped colonies. The colonies rapidly invert their curvature in response to changing light levels, which they detect through a rhodopsin-cGMP pathway. Inversion requires actomyosin-mediated apical contractility and allows alternation between feeding and swimming behavior. *C. flexa* thus rapidly converts sensory inputs directly into multicellular contractions. In this respect, it may inform reconstructions of hypothesized animal ancestors that existed before the evolution of specialized sensory and contractile cells.

**One Sentence Summary:** A newly described choanoflagellate species forms cup-shaped colonies that reversibly invert their curvature in response to light.

## Main Text

The evolution of animals from single-celled ancestors involved several major evolutionary innovations, including multicellularity, spatial cell differentiation, and morphogenesis (*1, 2*). Efforts to reconstruct the origin of animal multicellularity have benefited from the study of choanoflagellates, the closest living relatives of animals (*3–5*). Choanoflagellates are microbial eukaryotes that feed on bacteria and live in aquatic environments around the world; many species differentiate over their life history into diverse cell types, including unicellular and multicellular forms (*3, 6*–*8*). Comparative genomics and transcriptomics have revealed that many gene families once thought to be unique to animals (e.g. cadherins, C-type lectins, and receptor tyrosine kinases) are also present in choanoflagellates (*4, 9*–*12*). Moreover, laboratory studies of the model choanoflagellate *Salpingoeca rosetta* (*6*) have revealed diverse responses to environmental cues, such as pH-taxis (*13*), aerotaxis (*14*), and bacterial regulation of life history transitions (multicellular development (*15*) and mating (*16*)). However, *S. rosetta* is only one of approximately 380 known species (*17*) and choanoflagellates are at least as genetically diverse as animals (*9*). Choanoflagellate diversity thus represents a largely untapped opportunity to investigate environmental regulation of cell behavior, the principles that broadly underpin multicellularity, and the evolution of animal cell biology.

## Photic cues induce multicellular sheet inversion in a colonial choanoflagellate

During a survey of choanoflagellate diversity on the Caribbean island of Curaçao in April 2018 (Fig. 1, A and B), we collected large, cup-shaped colonies of protozoa (∼100 μm diameter) from shallow pools in the splash zone above the tide line of a rocky coastal area (Fig. 1B). Each colony was composed of a monolayer (“sheet”) of up to hundreds of flagellated cells (Fig. 1C, Movie S1). Upon closer inspection, we observed that the cells bore the characteristic collar complex of choanoflagellates (*1, 3*), in which a “collar” of microvilli surrounds a single apical flagellum (Fig. 1D). However, unlike in most choanoflagellate colonies (*3*), the apical flagella pointed into the interior of the colony (Fig. 1E), resembling the orientation of collar cells in the choanocyte chambers of sponges (*18*). While observing colonies, we were surprised to see them invert their curvature rapidly (within ∼30 s from initiation to completion) while maintaining their cell topology, such that the flagella now pointed outward along the radius of curvature of the colony (Fig. 1, F and G, Movie S2). The colonies tended to remain in the inverted form (“flagella-out”) for several minutes before reverting to their initial, relaxed conformation (“flagella-in”) in a similarly rapid process (Fig. 1H, Movie S3).

**Fig. 1.**
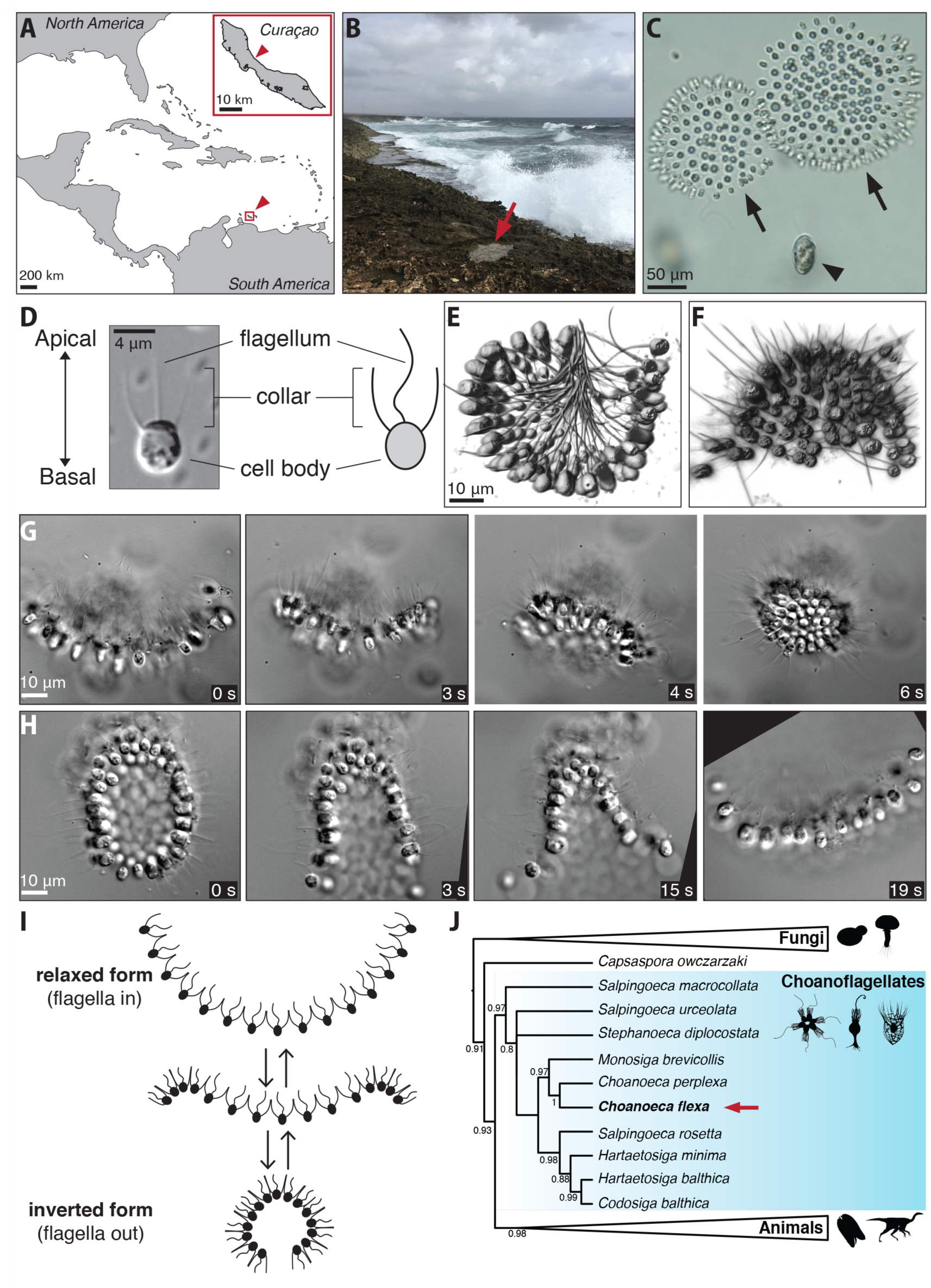
*Choanoeca flexa* sp. nov., a choanoflagellate discovered in splash pools on the island of Curaçao, forms colonies that rapidly and reversibly invert their curvature. (**A** to **C)** *Choanoeca flexa* was discovered in splash pools on the northern shore of Curaçao. (**A**) Map of the Caribbean Sea with arrowhead highlighting the island of Curaçao. Inset, magnified map of Curaçao with arrowhead indicating the sampling site (12°14’12.1” N, 69°01’34.8” W). (**B**) Photograph of the sampling site. Water samples were collected from splash pools (such as that indicated by the red arrow) within reach of ocean spray. (**C**) Light microscopy of freshly collected splash pool samples revealed a diverse microbial eukaryotic community, including dinoflagellates (*Oxyrrhis* sp.; arrowhead) and cup-shaped colonies of a previously unknown choanoflagellate (arrows), each comprising a monolayer of uniflagellate cells, that appeared to spontaneously invert their curvature (see Movie S1). The species name *flexa* was given in reference to this striking behavior. Shown is a still frame from Movie S1, recorded in Curaçao soon after sample collection. **(D)** *C. flexa* was recognizable as a choanoflagellate based on its cell morphology, visualized here by DIC microscopy. Each cell has a single apical flagellum surrounded by a collar of microvilli. **(E** and **F)** *C. flexa* colonies can adopt two distinct orientations, flagella-in and flagella-out, as visualized here through three-dimensional reconstruction of fixed colonies that were stained with a membrane marker (FM1-43FX). In the flagella-in orientation (**E**), the flagella project into the interior of the approximately hemispherical colony. In the flagella-out orientation (**F**), the flagella point outward from the hemisphere (same scale as E). **(G** and **H)** *C. flexa* colonies rapidly and reversibly invert their curvature while maintaining contacts among neighboring cells. (**G**) A flagella-in colony inverts to the flagella-out orientation and seals into a nearly closed sphere over the course of six seconds. Shown are still frames from Movie S2. (**H**) A flagella-out colony reverts to the flagella-in orientation through a similar process, as shown in still frames from Movie S3. Some frames have been rotated to facilitate tracking individual cells between images. **(I)** Summary schematic depicting the inversion process (in which sheets transit from flagella-in to flagella-out) and of the converse relaxation process (from flagella-out to flagella-in). **(J)** Phylogenetic analysis of 18S rDNA sequence confirmed that *Choanoeca flexa* (red arrow) is nested within the choanoflagellates and revealed that it is sister to the species *Choanoeca perplexa* (*19*). Fig. S1 shows the phylogeny with branch lengths to scale.

To start laboratory cultures of this choanoflagellate, we manually isolated several representative colonies away from the other microbial eukaryotes present in the original splash pool sample (e.g., presumptive *Oxyrrhis* sp. dinoflagellates; Fig. 1C, Movie S1) and transferred them into nutrient-supplemented artificial seawater along with co-isolated bacteria (which choanoflagellates need as a food source). Cells from the isolated colonies proliferated and served as the foundation of all downstream laboratory cultures. Phylogenetic analyses of 18S rDNA sequences indicated that it is the sister-species of the previously described species *Choanoeca perplexa* (also known as *Proterospongia choanojuncta*; Fig. 1I, Fig. S1), which has a dynamic life history that includes single cells and colonies (*7, 19*). In reference to the striking sheet bending behavior of the new species, we named it *Choanoeca flexa*. Interestingly, a similar inversion behavior was briefly mentioned in a 1983 study of *C. perplexa* (*7*). The cell line from that study subsequently stopped forming colonies in the laboratory and could never be revived nor re-isolated (*3*), preventing mechanistic study of the process. Our re-observation of this process in the newly discovered sister-species of *C. perplexa* confirms and extends that early report. This behavior, representing a global change in multicellular form, is reminiscent of concerted movement and morphogenesis in animals (e.g., muscle contraction or gastrulation). Because of the potential evolutionary implications of rapid shape change in *C. flexa*, we set out to investigate (1) how colony inversion is regulated, (2) the mechanisms underlying colony inversion, and (3) the ecological consequences of colony inversion.

Unexpectedly, several lines of evidence indicated that *C. flexa* colony inversion is regulated by light. While imaging live *C. flexa* sheets for long periods of time (>1hr) under constant illumination, we noted that colony inversions became less frequent. In contrast, after the microscope illumination was turned off, the colonies would invert almost immediately (Movie S7). To test whether light-to-dark transitions consistently induce *C. flexa* inversion, we established a quantitative assay based on the observation that the projected area of a *C. flexa* sheet decreases by as much as 50% during inversion (Fig. 2, A to C, Movie S4). Using this assay, we confirmed that a rapid decrease in illumination reliably induced inversion of *C. flexa* colonies from flagella-in to flagella-out within thirty seconds (Movie S5; Fig. 2D). Thus, *C. flexa* colony inversion can be triggered by light-to-dark transitions. To our knowledge, this represents the first observation of light-responsive behavior in a choanoflagellate.

**Fig. 2.**
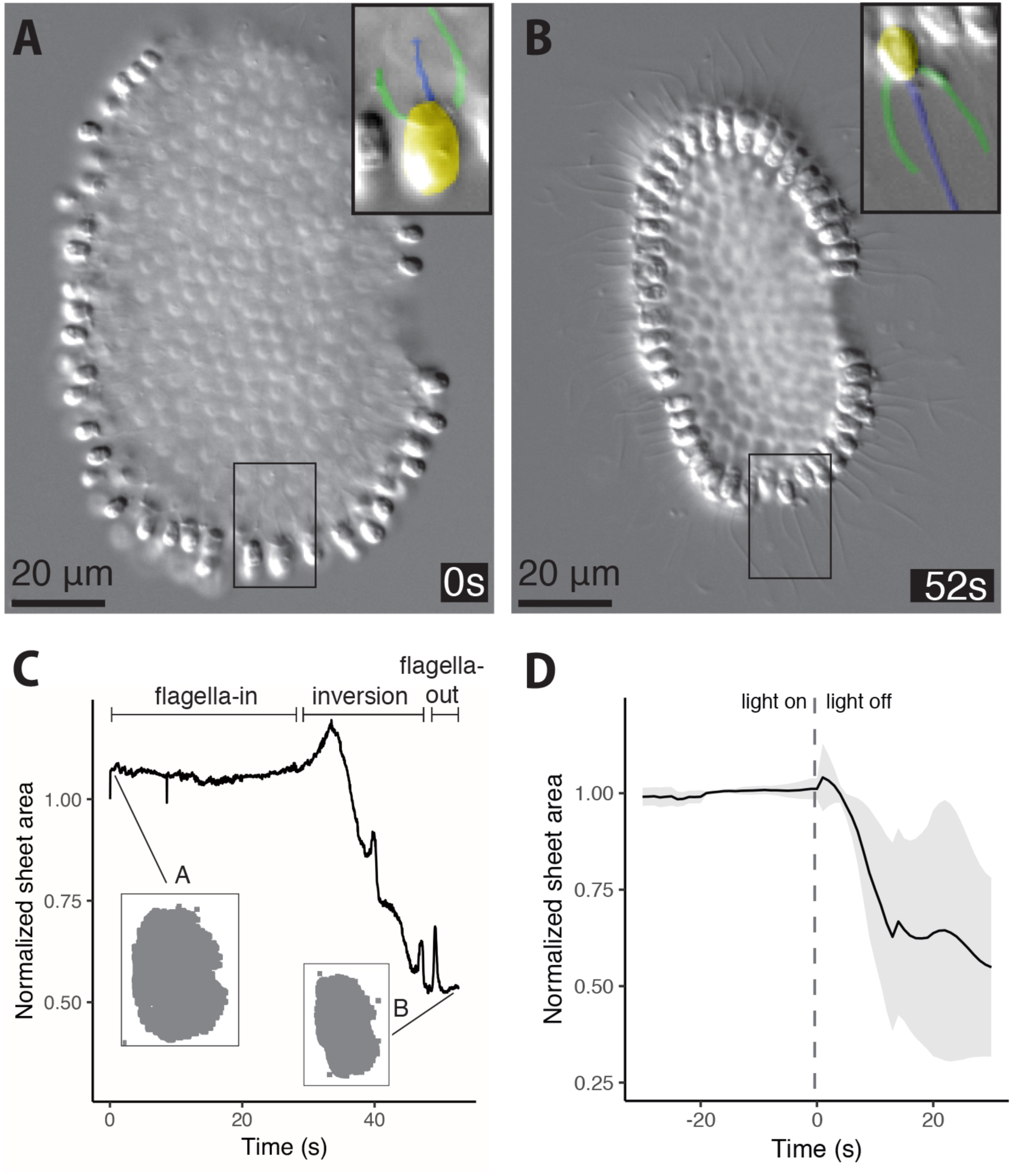
Light-to-dark transition induces *C. flexa* colony inversion. **(A** to **C)** Colony inversion can be tracked across a population at low magnification because it correlates with a decrease in the projected area of each colony. Time-lapse microscopy (Movie S4) shows a single *C. flexa* colony as it spontaneously inverted from the flagella-in (**A**) to flagella-out (**B**) orientation. Insets correspond to the boxed region of the colony, with pseudocoloring to highlight the orientation of the cell and its apical flagellum. Importantly, the orientation of the flagellum relative to the curvature of the colony inverts without the cell breaking its contacts with neighboring cells. (**C**) Projected surface area of the colony shown in panels (**A)** and (**B)** plotted as function of time. This inversion corresponded to a 50% decrease in area over ∼30 seconds. Normalized sheet area is defined as the area of a sheet divided by its initial value (at t = 0). **(D)** Colonies sense and respond to changes in light intensity, reliably undergoing inversion in response to light-to-dark transitions. Colony inversion can be quantified as a decrease in projected area. Area for *n =* 5 colonies, normalized by the initial value for each colony (at t=0), is plotted as a function of time before and after light reduction (vertical dotted line). See Movie S5 for a representative example. The line represents the mean projected area (rolling average over 5-second windows) and the ribbon represents standard deviation.

## A rhodopsin-cGMP pathway regulates colony inversion in response to light-to-dark transitions

We next sought to understand how *C. flexa* colonies detect and respond to a photic stimulus. Although choanoflagellates are unpigmented and transparent, at least four choanoflagellates (*9*) encode a choanoflagellate-specific rhodopsin-phosphodiesterase fusion protein (Fig. S2), RhoPDE, that has been investigated for its potential as an optogenetic tool (*20*–*23*). RhoPDE proteins consist of an N-terminal type I (bacterial) rhodopsin (a class of photosensitive transmembrane proteins broadly involved in light detection (*24*)) fused to a C-terminal phosphodiesterase (PDE) that catalyzes light-dependent cyclic nucleotide hydrolysis (Fig. 3A). Based on *in vitro* studies (*20–23*), RhoPDE from *S. rosetta* appears capable of converting a photic stimulus into a biochemical signal within seconds, similar to the time scale of the *C. flexa* response to light-to-dark transitions.

**Fig. 3.**
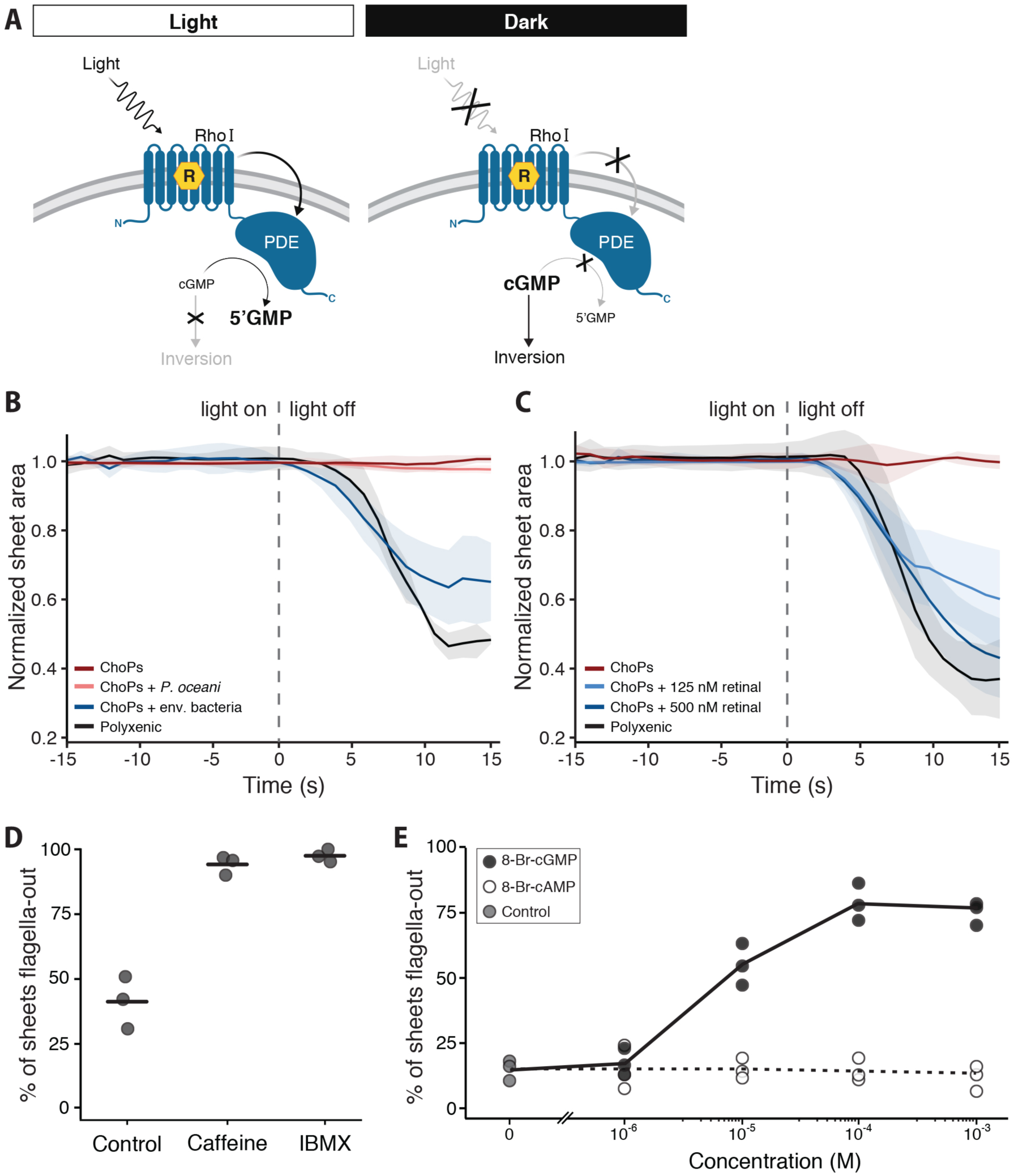
*C. flexa* cells transduce light stimuli through a rhodopsin-cGMP pathway using bacterial carotenoids. RhoPDE (blue), a choanoflagellate-specific enzyme rhodopsin and a candidate regulator of sheet inversion in response to light-to-dark transitions. *C. flexa* encodes four putative homologs of RhoPDE (Fig. S2), each comprising a type I (bacterial) rhodopsin (“RhoI”) fused to a cyclic nucleotide phosphodiesterase (“PDE”). Photodetection by rhodopsin requires the cofactor retinal (yellow hexagon, “R”), a covalently bound chromophore that undergoes isomerization in response to light (*28*). When light levels are high (left panel), the rhodopsin domain activates the PDE domain, which hydrolyzes cGMP to 5’GMP (*20–23*), keeping cellular cGMP levels low. When light is reduced (right panel), the PDE domain is inactive, allowing cellular cGMP levels to rise. We show below (**E**) that increased cellular cGMP leads to inversion. **(B)** A bacterially produced factor is required for light-regulated sheet inversion. *C. flexa* sheets were grown in the presence of different combinations of bacteria and their photic response was quantified as in Fig. 2D. *C. flexa* sheets grown in a “Polyxenic” culture containing diverse co-isolated environmental bacteria (Table S1) inverted to flagella-out in response to decreased illumination, as measured by a decrease in colony projected area. By contrast, sheets grown in a monoxenic culture that contains only *C. flexa* and the bacterium *P. oceani* (“ChoPs” culture), did not respond to changes in illumination. When the ChoPs culture was inoculated with environmental bacteria from the polyxenic culture (“ChoPs + env. bacteria”), the photic response was restored, showing that a bacterial factor is necessary for light-regulated inversion. As a negative control, ChoPs inoculated with only *P. oceani* bacteria (“ChoPs + *P. oceani*”) did not show restoration of the inversion response. Shown are data from *n =* 4 ChoPs colonies, *n = 3* Polyxenic colonies, *n =* 3 colonies of ChoPs + env. bacteria, and *n =* 3 colonies of ChoPs + *P. oceani*. **(C)** The rhodopsin chromophore retinal (or its carotenoid precursors) is the bacterial molecule required for the photic response. The ChoPs culture, which is insensitive to changes in illumination, was treated with varying concentrations of retinal, and the photic response was quantified as in Fig. 2D. Treatment with 125 nM or 500 nM retinal was sufficient to restore light-regulated inversion. Thus, the light-insensitive phenotype of the ChoPs strain is due to the absence of a bacterial carotenoid, which indicates that rhodopsin activity is required for inversion in response to light-to-dark transitions. Shown are data from *n =* 4 ChoPs colonies, *n = 4* Polyxenic colonies, *n =* 5 colonies of ChoPs + 125 nM retinal, and *n =* 5 colonies of ChoPs + 500 nM retinal. **(D)** Phosphodiesterase activity suppresses sheet inversion in *C. flexa*. Treatment with the phosphodiesterase inhibitors caffeine (10 mM) or IBMX (1 mM) caused *C. flexa* colonies to invert to the flagella-out orientation in the absence of a photic stimulus (*n* = 3 independent trials; *N* = 52, 55, and 38 for controls; *N* = 23, 31, and 40 for caffeine; *N* = 42, 37, and 27 for IBMX). **(E)** cGMP acts as a second messenger in the *C. flexa* phototransduction pathway. Treating the light-unresponsive ChoPs culture with increasing concentrations of a cell-permeant cGMP analog (8-Br-cGMP) caused sheets to invert into the flagella-out orientation in a dose-dependent manner in the absence of a photic stimulus. By contrast, treating with 8-Br-cAMP did not have an effect. Thus, increased cellular cGMP concentration is sufficient to cause sheet inversion.

To test for the presence of RhoPDE or other candidate photosensitive proteins in *C. flexa*, we sequenced and assembled the *C. flexa* transcriptome (Figshare DOI 10.6084/m9.figshare.8216291). 56 million reads were assembled into 50,463 predicted transcripts encoding 20,477 predicted unique proteins, of which four appeared to be RhoPDE homologs (Fig. S2; GenBank accession numbers MN013138, MN013139, MN013140 and MN013141). No other rhodopsins were detected in the *C. flexa* transcriptome. The only other candidate photoreceptor protein domain found was a member of the cryptochrome family of photosensitive transcription factors (*25*), which act on the timescale of transcriptional regulation (at least several minutes (*26, 27*)) and therefore appear unlikely to mediate the light-to-dark transition response. Thus, we focused our attention on the RhoPDEs.

The hypothesized role of RhoPDE as the regulator of the light-to-dark transition response offered two testable predictions. First, depletion of the rhodopsin chromophore, retinal (*28*), should prevent the response by abolishing rhodopsin activity. Second, artificially increasing the cellular concentration of cGMP or cAMP (which are degraded by the enzymatic activity of PDEs) should mimic the effect of darkness and therefore be sufficient to trigger sheet inversion.

Retinal is a carotenoid chromophore whose isomerization underlies rhodopsin photodetection (*28–30*). Plants and some bacteria can synthesize carotenoids, but transcriptome analysis revealed that *C. flexa* lacks a key enzyme in the retinal biosynthesis pathway (Fig. S3); therefore, like animals, *C. flexa* must receive retinal or its biochemical precursor, beta-carotene, from its food. Because *C. flexa* sheets are grown with diverse co-isolated environmental bacteria, it is possible that they take up retinal or beta-carotene from their bacterial prey. To test whether bacterially produced carotenoids are required for light-regulated colony inversion, we established a monoxenic culture containing only *C. flexa* and a co-isolated bacterial species, *Pseudomonas oceani*, that lacks genes in the retinal biosynthesis pathway (Fig. S3) (*31*). This culture, referred to as “ChoPs” (for *<u>Cho</u>anoeca* + *<u>Ps</u>eudomonas*), was expected to be devoid of carotenoids, thereby abolishing rhodopsin activity. As predicted, *C. flexa* sheets in ChoPs cultures did not invert in response to darkness (Fig. 3B). Inoculating ChoPs cultures with a mixture of additional co-isolated environmental bacteria restored the light-to-dark response, demonstrating that a bacterial factor is necessary for this behavior. Moreover, addition of exogenous retinal to ChoPs cultures was sufficient to restore the wild-type light-to-dark response in *C. flexa* (Fig. 3C). The requirement of retinal for the inversion response and the fact that the only rhodopsin-containing genes in the *C. flexa* genome encode RhoPDEs suggest that one (or more) RhoPDEs are required for the light-to-dark-induced inversion.

We next investigated whether cyclic nucleotide signaling plays a role in *C. flexa* phototransduction. Treatment of *C. flexa* sheets with two inhibitors of phosphodiesterase activity, caffeine (*32*) and IBMX (*33*), caused colonies to invert even in the absence of a photic stimulus (Fig. 3D). Moreover, incubating *C. flexa* sheets with a cell-permeant analog of cGMP induced colony inversion in a dose-responsive manner, while a cell-permeant analog of cAMP had no effect (Fig. 3E), suggesting that cGMP acts as a second messenger in phototransduction and is the endogenous trigger of colony inversion.

Together, these results provide evidence that the *C. flexa* response to light-to-dark transitions relies on a rhodopsin as a photoreceptor and on cGMP as a second messenger. The simplest interpretation of these findings is that a RhoPDE protein controls *C. flexa* phototransduction. However, direct validation of this hypothesis will require targeted disruption of the RhoPDE homologs encoded by *C. flexa* (Fig. S2), which is not currently possible.

## Sheet inversion mediates a trade-off between feeding and swimming

What are the functional and ecological roles of sheet inversion in *C. flexa*? Flagella-in sheets showed little to no motility, either slowly drifting or settling to the bottom of culture flasks (Movie S6). In contrast, inverted (flagella-out) sheets swam rapidly (Fig. 4, A and B, Fig. S5, Movie S7). Thus, one important consequence of sheet inversion is increased motility, which could allow rapid escape from environmental hazards (including predators).

**Fig. 4.**
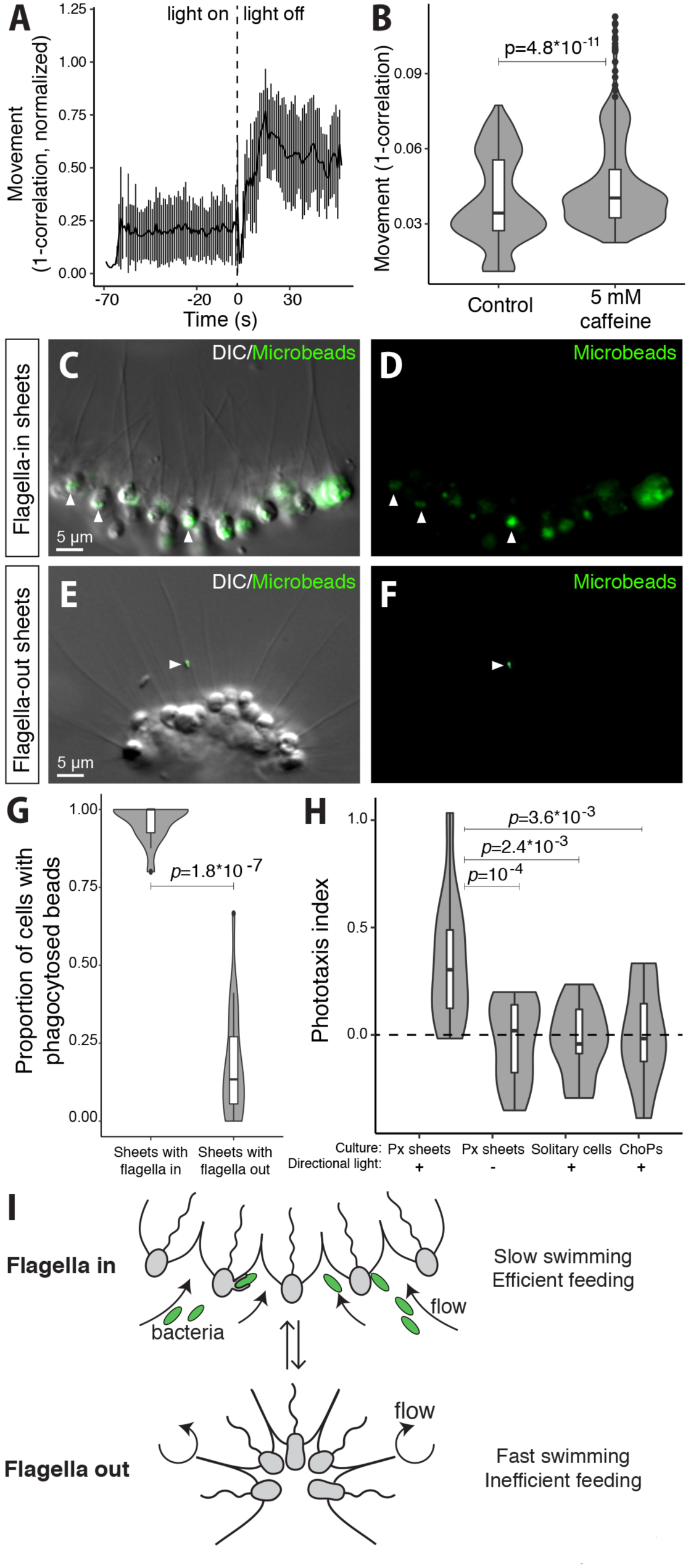
Sheet inversion results in a trade-off between swimming and feeding. **(A** and **B)** Flagella-out sheets swim faster than flagella-in sheets. **(A)** Following light-to-dark induced inversion, flagella-out sheets swam faster than they did in their relaxed, flagella-in form. Swimming speed increased quickly after darkness-induced inversion (Movie S7), as quantified by an increase in the measured amount of movement. Movement was defined as 1-correlation (where “correlation” refers to the Pearson correlation between two consecutive frames of a given movie; see Material and Methods). Movement was normalized between 0 and 1 for each of *n* = 9 time-lapse movies. Error bars represent standard deviation. **(B)** Sheets swim faster after caffeine-induced inversion. Here, caffeine treatment (5 mM) was used to chemically induce inversion in all sheets under constant light, presumably by inhibiting PDE activity. Caffeine treatment thus enabled sustained experiments with inverted sheets by preventing relaxation. Movement was quantified as in (**A**). *n =* 9 time-lapse movies for the control condition (populations of relaxed sheets imaged under constant ambient light) and *n* = 10 movies of sheet populations in which inversion has been induced by 5 mM caffeine. **(C** to **G)** Flagella-in sheets feed more efficiently than flagella-out sheets. Choanoflagellates feed by phagocytosis of bacterial prey captured from the water column. Internalization of 0.2 μm fluorescent beads was used to quantify phagocytic activity of cells in sheets. (**C** to **F**) Detection of beads phagocytosed by flagella-in sheets (untreated) and flagella-out sheets (treated with 5 mM caffeine) that were fixed after incubation for 1 hour with fluorescent microbeads. Cells were visualized by DIC transmitted light (**C** and **E**) and beads by green fluorescence **(C** to **F)**. Arrowheads: fluorescent beads (inside the cells in C, stuck to a flagellum in E). **(G)** Proportion of cells having phagocytosed beads in *n =* 17 sheets with flagella in compared to *n =* 21 sheets with flagella out. *p*-value is by the chi-square test. **(H)** Sheet phototaxis requires retinal and multicellularity. Polyxenic sheets (capable of perceiving light) migrate toward a lateral light source over 1 hour (*n* = 12 experiments). By contrast, no directional accumulation was observed in polyxenic sheets without directional light (*n* = 12 experiments). Interestingly, dissociated single cells (*n* = 9 experiments) were not capable of phototaxis. Likewise, retinal-deprived monoxenic cultures (ChoPs) that do not invert in response to light-to-dark transitions are also incapable of phototaxis (*n* = 10 experiments). *P*-values are by an ANOVA with Dunnett’s correction. **(I)** Model of how sheet inversion mediates a swimming-feeding tradeoff. In flagella-in sheets, flagellar beating generates a feeding flow that carries bacteria toward the basal side of the cells (Fig. S7), allowing phagocytosis. In flagella-out sheets, flagellar beating allows swimming, and the basal side of the cells faces the inner side of the colony, preventing it from coming in contact with bacterial prey.

In choanoflagellates, flagellar beating in unattached, single cells typically results in motility, while flagellar beating in cells attached to surfaces or other cells (e.g. in colonies) has been hypothesized to enhance feeding currents that draw bacterial prey to the outside of the collars for phagocytosis (*3, 34*). Hence, the enhanced motility of flagella-out colonies might come at a cost – reduced feeding efficiency. To test for the existence of a tradeoff between swimming and feeding, we used bacteria-sized fluorescent beads (*35*) to quantify particles ingested by cells of flagella-in and flagella-out sheets (Fig. 4, C to F). While cells in flagella-in sheets fed efficiently (>75% cells/sheet internalizing beads, Fig. 4G), cells in flagella-out sheets did not (∼10% cells/sheet internalizing beads on average, Fig. 4G). Moreover, the flow generated by the sheets during inversion was visualized by tracking fluorescent beads in suspension in sea water. In relaxed sheets, the flow converged toward the center of the colony (carrying bacterial prey toward the cells), while in inverted sheets, the flow was directed away from the colony – allowing swimming, but not feeding (Fig. S6).

Because sheets swim slowly when relaxed (flagella-in) and rapidly when inverted (flagella-out), we suspected that darkness-induced inversion might allow sheets to accumulate in bright areas, effectively undergoing phototaxis. To test for phototaxis, we illuminated chambers containing *C. flexa* sheets with directional light and found that sheets tended to accumulate in the brightest areas near the illumination source compared to a control in which no illumination was provided (Fig. 4H, Fig. S7). Further, we found that neither colonies from ChoPs cultures nor single cells from dissociated sheets were capable of phototaxis, suggesting that rhodopsin activity, multicellularity, and sheet inversion are all required for phototaxis (Fig. 4H). These results demonstrate that sheet inversion mediates a tradeoff between feeding (flagella-in) and swimming (flagella-out) that is plausibly ecologically relevant (Fig. 4I).

## Sheet inversion requires apical actomyosin contractility

How do cells in sheets interact with their neighbors and what cellular mechanisms allow sheet inversion? By employing DIC microscopy on live colonies, as well as confocal microscopy and electron microscopy on fixed colonies, we found that cells in *C. flexa* sheets are linked by direct contacts between collar microvilli (Fig. 5, A to C; Fig. S8). Importantly, we found no evidence for the intercellular bridges, shared ECM, or filopodial contacts that mediate multicellularity in other choanoflagellate species (*6, 7, 10, 36*).

**Fig. 5.**
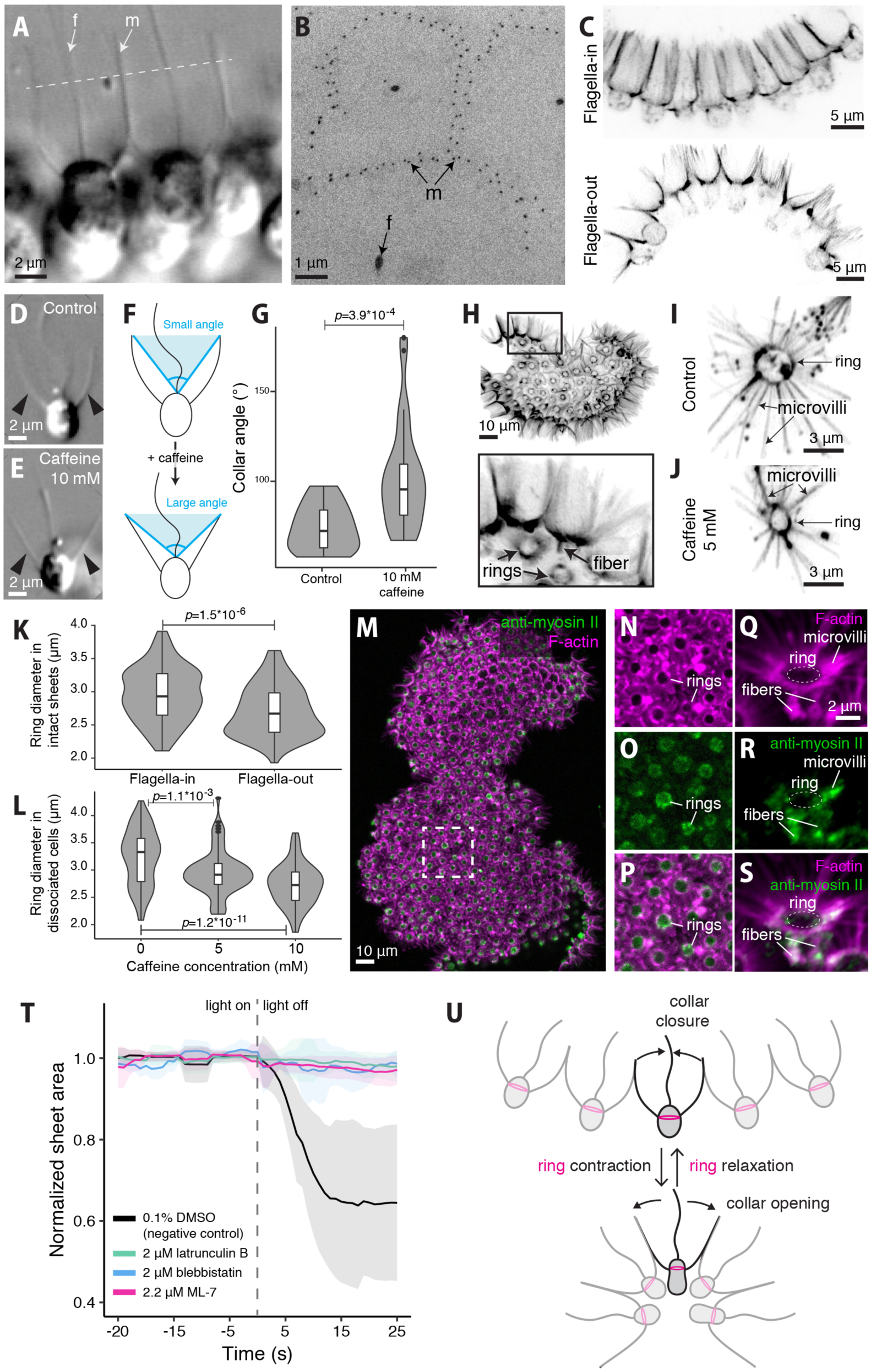
Sheet inversion requires apical actomyosin cell contraction. **(A** and **B)** Cells within a sheet are linked by direct contacts between their collars. (A) Transmitted light DIC microscopy image, showing direct contact between the microvillous collars of neighboring cells. (m): microvilli, (f): flagellum. Dotted line indicates approximate plane of transverse section in panel **B. (B)** Transmission electron microscopy of a transverse section through the microvillar collars of neighboring cells reveals close contacts microvilli. **(C)** Collar morphology differs between flagella-in sheets (top) and flagella-out sheets (bottom). Cells in flagella-in sheets have collars that are cylindrical (with the microvilli nearly parallel) while the microvilli of flagella-out sheets are flared, producing conical collars. Collars were stained with fluorescent phalloidin in fixed samples and imaged by confocal microscopy. Both panels are maximum projections of Z-stacks. **(D** to **G)** Caffeine treatment of dissociated cells caused the collar to flare out, resembling the collars found in flagella-out sheets. (**D** and **E**) Two representative individual cells imaged by DIC microscopy, either without (**D**) or with (**E**) caffeine treatment. Arrowheads: microvilli. (**F**) The differences in collar morphology where quantified in terms of the collar angle (defined by the tip of two bilateral microvilli and the base of the flagellum). (**G**) Collar angles in caffeine-treated cells are significantly larger than those in control cells. Collar angles measured in *n* = 16 untreated dissociated cells and *n* = 28 caffeine-treated cells. Data are presented as violin boxplots, showing the median value (black horizontal lines inside white boxes), interquartile range (white boxes), and total range (thin lines). Surrounding the boxplots are kernel density traces plotted symmetrically (violin plots). *p*-value by Mann-Whitney’s U test. **(H)** An actin ring connected to a small number of longitudinal fibers is present at the base of each collar, visualized here in a sheet stained with fluorescent phalloidin and imaged by confocal microscopy (maximum projection of a Z-stack). Inset: higher magnification shows actin rings and a longitudinal fiber extending below collar microvilli. **(I** to **L)** The actin ring constricts during inversion in intact sheets and in response to caffeine in isolated cells. (**I** and **J**) Actin ring observed by phalloidin staining in an untreated cell **(I)** and in a cell treated with 5 mM caffeine (**J**). (**K**) Ring diameter is consistently larger in flagella-in sheets (*n =* 8 sheets, N = 124 cells) than in flagella-out sheets (*n* = 7 sheets, N = 110 cells). *p =* 1.52*10-6 by Mann-Whitney’s U test. (**L**) Ring diameter is consistently smaller in 10 mM caffeine-treated dissociated cells (*n =* 74) and 5 mM caffeine-treated dissociated cells (*n* = 82) than in untreated dissociated cells (*n* = 89). *p-*values by Dunnett’s test for comparing several treatments with a control. **(M** to **S)** Antibodies against myosin II localize to the apical actin ring, to the longitudinal fibers, and to the base of microvilli in immunostained sheets. (**M**) Immunostained sheet, (**N** to **P**) close-up views showing the apical ring, (**Q** to **S**) close-up view showing the fibers and base of microvilli. Green: Sigma M7648 rabbit anti-myosin II antibody, magenta: rhodamine-phalloidin. (Also see Fig. S10.) **(T)** Treatment with inhibitors of actin polymerization (latrunculin B, *n* = 6 colonies), myosin contractility (blebbistatin, *n* = 6 colonies), or myosin activation by phosphorylation (ML-7, *n* = 9 colonies) prevented sheet inversion in response to light-to-dark transitions. *n* = 13 colonies for the DMSO-treated (negative control) condition. Photic response was quantified as in Fig. 2D. **(U)** Summary schematic of the proposed inversion mechanism. Contraction of the apical actomyosin network (comprising ring, fibers, and base of the microvilli) correlates with, and is required for, collar flaring and sheet inversion.

Interestingly, we also observed that collar morphology tended to differ between relaxed and inverted sheets. In relaxed sheets (flagella-in; Fig. 5C), the microvilli on each cell assembled into a barrel-shaped collar whose diameter varied little from base to tip. In inverted sheets (flagella-out; Fig. 5C), the microvilli formed a flared, cone-shaped collar whose diameter increased from base to tip. This contrast in collar shape suggests a potential mechanism for sheet inversion: active “opening out” of the collar, by increasing the surface area of the apical side of the sheets relative to their basal side, might force a change in sheet curvature. Consistent with this, we found that, as in intact sheets, dissociated *C. flexa* cells treated with caffeine opened their collar into a conical shape, while untreated controls maintained a barrel-shaped collar (Fig. 5, D to G). Additionally, caffeine treatment caused the microvilli to straighten and the base of the collar to slide toward the equator of the cell (Fig. S9). These data indicate that the changes in collar geometry observed during inversion are actively generated by individual cells, and do not require interactions among neighboring cells.

If *C. flexa* cells modulate the shape of their collars, what is the underlying cellular mechanism? In animal epithelial tissues, sheet bending during morphogenesis is frequently due to a process called apical constriction, in which contraction of an apical actomyosin network reduces the surface area of the apical side of the cell (*37, 38*). Apical constriction is mediated by molecular motors belonging to the myosin II family, which is of ancient eukaryotic origin (*39*) and is represented in all previously sequenced choanoflagellate genomes (*40*) and transcriptomes (*9*). The *C. flexa* transcriptome encodes homologs of the myosin II regulatory light chain (GenBank accession number MK787241) and heavy chain (GenBank accession number MK787240) (Fig. S4) whose protein sequences are respectively 78% and 63% similar to their human counterparts.

We thus investigated the actomyosin cytoskeleton of *C. flexa*. Like in choanoflagellates, the apical side of animal epithelial cells is defined by the presence of a cilium/flagellum and/or microvilli, and the apicobasal axis of both types of cells is broadly accepted to be homologous (*41*). Confocal imaging of sheets labelled with fluorescent phalloidin revealed the presence of a pronounced F-actin ring at the apical pole of each cell, from which the microvillar collar extends (Fig. 5H). Directly connected to this ring, and perpendicular to it, we detected a small number of longitudinal actin fibers (usually two or three) pointing toward the basal pole. Diameter measurements showed that the actin ring was consistently smaller in inverted, flagella-out sheets compared with relaxed, flagella-in sheets (Fig. 5, I to L). The same was true for dissociated, caffeine-treated cells compared with the corresponding negative controls (Fig. 5, I-J and L), consistent with the ring actively constricting during sheet inversion. During inversion and in response to caffeine, some (but not all) cells transiently acquired a “bottle cell” morphology with a narrow apex and a bulbous base (Fig. S9), reminiscent of animal cells undergoing pronounced apical constriction (*42, 43*). Interestingly, caffeine treatment also induced shortening of the longitudinal actin fibers (Fig. S9, G to H), suggesting that fiber contraction pulls the collar toward the basal pole.

Using five different commercial myosin II antibodies, including two raised against the activated phosphorylated form (Fig. S10), we found that *C. flexa* cells contain myosin that overlaps in regions with the apical actin ring (Fig. 5, M to P, Fig. S10), longitudinal fibers, and base of the microvilli (Fig. 5Q-S), consistent with the idea that the apical actin network is contractile. Finally, blebbistatin (which inhibits the ATPase activity of myosin II (*44*)) entirely abolished ring constriction in caffeine-treated dissociated cells (Fig. S11) and prevented sheet inversion (Fig. 5T), as did latrunculin B (which inhibits dynamic actin polymerization (*45*)) and ML-7 (which prevents activation of myosin by phosphorylation of the Myosin Regulatory Light Chain (*46*)) (Fig. 5T). None of these drugs affected flagellar beating (which was used as a visual control of cell survival, see Material and Methods), consistent with them specifically targeting actomyosin. Together, these results suggest that sheet inversion requires apical constriction of an actomyosin network at the base of the collar (Fig. 5U).

## The ancestry of apical constriction

The discovery of sheet bending driven by apical constriction in a multicellular choanoflagellate has several potentially important evolutionary implications. Epithelial sheet bending is a fundamental mechanism underlying animal embryonic development (*37, 47, 48*) and multicellular contractility also plays a fundamental role in the behavior of adult animals by allowing fine-tuned body deformations (*49*). As both embryonic and adult tissue contractility are found in nearly all animal lineages, including sponges (*50, 51*), ctenophores (*52, 53*), placozoans (*54*), cnidarians (*42, 55*) and bilaterians (*37, 38, 56*), both were likely present in the last common animal ancestor. By contrast, collective contractility and apical constriction were hitherto unknown in close relatives of animals, making their origin mysterious.

The existence of actomyosin-mediated apical constriction in *C. flexa* raises the possibility that this cellular module might have been present in the last common ancestor of choanoflagellates and animals (which together comprise the choanozoans (*1*)). Collar contractions have been reported in the unicellular stages of three other choanoflagellate species: *Codosiga pulcherrima* (*57*), *Monosiga gracilis* (*58*), and the sessile form of *C. perplexa* (*7*), the sister-species of *C. flexa*. As part of this study, we observed collar contractions in *C. flexa* sheets (Fig. 5G), dissociated cells from sheets (Fig. 5, D and E), and in naturally solitary “thecate” cells (Movie S12). We expanded this taxonomic sampling by investigating four other choanoflagellate species, *Monosiga brevicollis, S. rosetta, Salpingoeca urceolata* and *Diaphanoeca grandis*, which together cover the three main branches of the choanoflagellate phylogenetic tree (*17*). All four species displayed spontaneous changes in collar geometry occurring at the scale of a few seconds. *S. urceolata* (Movie S8) and *M. brevicollis* (Movie S9) showed spontaneous and reversible opening/closing of the collar (similar to *C. flexa*), while *S. rosetta* (Movie S8) and *D. grandis* (Movie S11) displayed subtler shape changes (reorientation of individual microvilli and modulation of collar curvature, respectively) (Figure 6A; Fig. S12). In all four species, immunostaining revealed the presence of an apical actomyosin ring at the base of the collar. This suggests that the apical actomyosin ring is a conserved feature of choanoflagellate biology (Figure 6, B and C) and that unicellular apical constriction was present in the last common ancestor of choanoflagellates and animals.

**Fig. 6.**
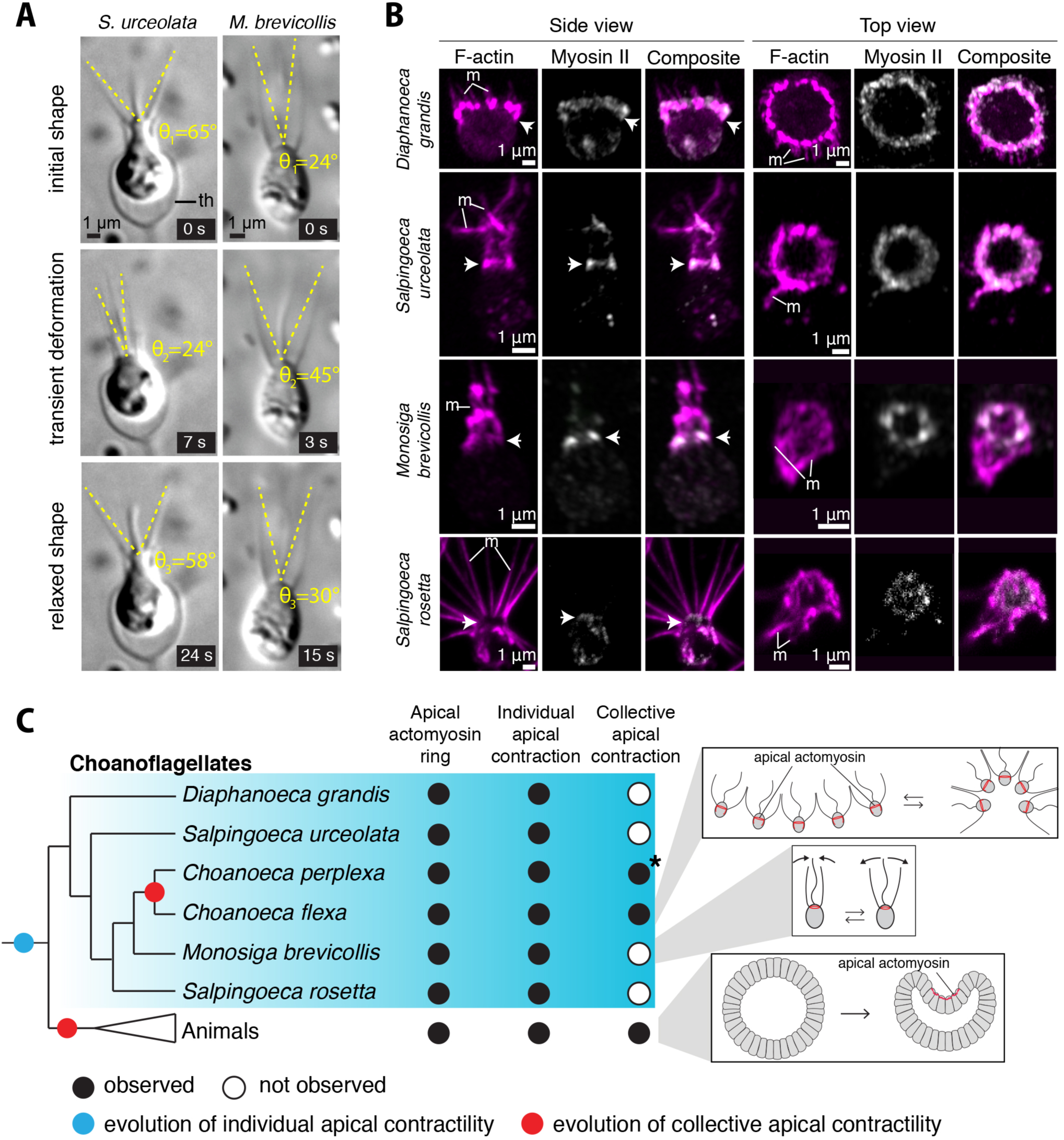
Apical constriction is conserved in choanoflagellates. **(A)** Spontaneous collar contractions observed in *Salpingoeca urceolata* and *Monosiga brevicollis* by time-lapse DIC microscopy. Dark and gray traces represent cell outline before and after contraction (3 to 7 seconds later), respectively. *S. urceolata* shows global and reversible collar closure correlated with retraction of the cell within its theca (Movie S8). *M. brevicollis* shows transient and reversible opening of its collar (Movie S9). (th): theca. **(B)** An apical actomyosin ring is detected at the base of the collar in four different choanoflagellate species: *Diaphanoeca grandis, S. urceolata, M. brevicollis* and *Salpingoeca rosetta*. F-actin stained by rhodamine-phalloidin. Myosin II stained with the Sigma M7648 antibody. Composite shows overlay of F-actin and myosin II staining patterns. Note that myosin II was generally not detected in the microvilli, except in *S. urceolata*. (m): microvilli **(C)** Apical constriction of individual cells was present in the last common ancestor of choanoflagellates and animals, and independently gave rise to multicellular apical constriction in *C. flexa* and in animals (see Figs. 6A and S13 for supporting data.). \**C. perplexa*, the sister-species of *C. flexa*, can undergo transient inversions of colony curvature that were briefly reported in an earlier study (*7*). Based on our study of *C. flexa*, we hypothesize that these inversions reflect conservation of collective apical constriction with that observed *C. perplexa*. Unfortunately, the currently available *C. perplexa* strains (from frozen stocks stored in the ATCC and in our lab) no longer form colonies in culture.

What might be the function of apical constriction in single cells? In some sessile choanoflagellates – including in the thecate form of *C. perplexa* – collar contraction happens in response to physical contact with an external object, and allows retraction of the cell inside an extracellular structure (called a theca) (*19*), suggesting it represents a defensive withdrawal reflex from predators or other threats. In free-swimming cells, collar contraction might fine-tune the hydrodynamics of swimming and/or feeding: for example, a closed collar might reduce drag and facilitate locomotion, while a spread collar could slow down swimming and increase collar area, thereby facilitating prey capture. Validation of these functional hypotheses will require direct testing.

These observations suggest that apical actomyosin mediated cell constriction evolved on the choanozoan stem lineage (Figure 6B). Could this cellular module be even more ancient? Polarized actomyosin contractions have been implicated in multicellular morphogenesis in the fruiting body of the slime mold *Dictyostelium* (*59*) and may be homologous to those observed in choanoflagellates and animals. However, the absence of comparable processes in the intermediate branches between *Dictyostelium* and choanozoans raises the possibility that polarized cell contractions in *Dictyostelium* and apical constriction in choanozoans evolved independently (*60*). Finally, the ichthyosporean *Sphaeroforma arctica*, a close relative of choanozoans, forms large multinucleated spores that partition into distinct cells in an mactomyosin-dependent process (*61*), providing an independent example of actomyosin-dependent multicellular development.

In contrast to single-cell apical constriction, the multicellular sheet bending observed in *C. flexa* and *C. perplexa* (*7*) has not been reported in other choanoflagellates. This suggests that apical constriction was present in solitary cells in the last choanozoan common ancestor, and was independently converted into multicellular sheet bending through the evolution of intercellular junctions in animals (*62*) and the evolution of microvillar adhesions in *C. flexa*. Interestingly, multicellular inversion has been proposed to have been part of the developmental repertoire of ancient animals (*1, 63*), based on the existence of whole-embryo inversion (from flagella-in to flagella-out) during calcareous sponge development (*64*). It is also intriguing that a similar inversion (but much slower – about an hour-long) takes place during the development of the alga *Volvox* (*65*). Given the large evolutionary distance between choanoflagellates and volvocalean green algae, along with the absence of inversion in intervening branches, inversion likely evolved independently in both groups (*1*).

In animals, the control of multicellular contractions invariably relies either on the cooperation of multiple cell types (as in adult organisms (*55, 56, 66*)) or on complex programmed signaling cascades (as in embryos (*37, 47, 48*)). By contrast, *C. flexa* directly converts sensory stimuli into collective contractions, without observable spatial cell differentiation, and evokes some hypotheses of early animal evolution that envisioned the first contractile tissues as homogeneous myoepithelia of multifunctional sensory-contractile cells (*67*).

The fact that contractility in *C. flexa* can be controlled by light represents another intriguing parallel to animal biology. Indeed, rhodopsin-cGMP pathways similar to that in *C. flexa* also underlie phototransduction in some animal cells (e.g. bilaterian ciliary photoreceptors (*29, 30*) and cnidarian photoreceptors (*68, 69*)), as well as in fungal zoospores (*70*). In contrast with choanoflagellates, however, phototransduction in animal photoreceptors relies on a type II (eukaryotic) rhodopsin that activates a separate phosphodiesterase through a G-protein intermediary (*29, 30*) (Fig. S13). Meanwhile, fungal zoospores use a distinct rhodopsin fusion protein (a type I rhodopsin fused to a guanylyl cyclase) to increase cellular cGMP in response to light (*70*) (Fig. S13). If a RhoPDE fusion protein controls *C. flexa* phototransduction, this would represent a third unique solution to the problem of transducing information from a change in illumination into a change in cyclic nucleotide signaling.

Much remains to be discovered concerning the ecological function, mechanical underpinnings, and molecular mechanisms of phototransduction and apical constriction in *C. flexa*. A deeper understanding will require the development of molecular genetic tools, which have only recently been established in *S. rosetta* (*36, 71*) and *D. grandis* (*72*). Nonetheless, *C. flexa* demonstrates how the exploration of choanoflagellate diversity can reveal unexpected biological phenomena and provides an experimentally tractable model for studying multicellular sensory-contractile coupling.

## Acknowledgments

We thank the Caribbean Research and Management of Biodiversity Foundation (CARMABI) for hosting us during field work, the Canadian Institute for Advance Research for their support of field work in Curaçao, and Patrick Keeling and his team for access to the microscope used to generate figure 1C. We thank the staff and students of the 2018 Physiology course at the MBL in Woods Hole, Zeiss for access to an AxioZoom, and Therese Gerbich and Tanner Fadero for help and advice on early phototaxis experiments. We thank Danielle Jorgens and Guangwei Min (from the Electron Microscopy Laboratory at UC Berkeley) for help with SEM. We thank Maxwell Coyle and Christian Erikson for advice on transcriptome assembly. We thank Monika Sigg and the Berkeley Functional Genomic Laboratory for help with iTag sequencing. Finally, we thank David Bilder, Marla Feller, Bob Goldstein and Kristin Scott for critical feedback on the manuscript, and the whole King lab for stimulating discussions.

## Author contributions

TB, BTL, and TAL: conceptualization, investigation, methodology, formal analysis, visualization, writing. MJAV: project administration and resources. KLM: investigation and visualization. NK: conceptualization, supervision, project administration and writing.

## Data and materials availability

Raw RNAseq reads used to assemble the *C. flexa* transcriptome have been deposited at the NCBI SRA with the accession number PRJNA540068 under BioProject PRJNA540068 and BioSample SAMN11533889. Transcriptome and predicted non-redundant proteome are available on Figshare at DOI: <u>10.6084/m9.figshare.8216291</u>. mRNA sequences and predicted amino acid sequences for genes of interest have been deposited on GenBank: myosin regulatory light chain (MK787241), myosin heavy chain (MK787240), and the four rhodopsin-phosphodiesterase paralogs (MN013138, MN013139, MN013140 and MN013141).

