## Supplementary Materials for "Light-regulated collective contractility in a multicellular choanoflagellate"

**This PDF file includes:**

Materials and Methods……………………………………………2
Figs. S1 to S13……………………………………………………..12
Table S1……………………………………………………………31
Captions for Movies S1 to S12…………………………………….32

**Other Supplementary Materials for this manuscript include the following:**

Movies S1 to S12

### Materials and Methods

#### Species description

Order Craspedida Cavalier-Smith 1997 (*73*).
Genus *Choanoeca* Ellis 1930 (*74*)
*Choanoeca flexa* sp. nov.

**Etymology**: from the Latin 'flexa', nominative feminine singular of 'flexus' which means 'bending' or 'transition, change'.
**Type locality**: splash pool of the Curaçao rocky coast (12.22831° N, -69.01353° W), from which water was collected with 25 $cm^2$ canted U-shaped culture flasks (T25; ThermoFischer Scientific 1012628) in April 2018.
**Description:** cell body is about 4-5 μm long. Microvillous collar is about 10 μm long. Microvilli of resting cells appear markedly curved. Cells form two-dimensional curved sheets linked by direct contact between their collars. Resting sheets are cup-shaped hemispheres with flagella pointing on the inner side of the curvature. Cells in their thecate form (Movie S12) are characterized by a stalkless theca, a bulbous cell base, a narrow and pointed apex, and a widely spread collar without a flagellum, as in the thecate form of *Choanoeca perplexa* (*19*).
**Holotype**: Figure 1E.

#### Initial isolation and culture of *Choanoeca flexa*

Samples containing *Choanoeca flexa* were first isolated by the three co-first authors from multiple splash pools on the northern rocky coast of Curaçao (12°14'12.1" N, 69°01'34.8" W). Individual colonies were manually isolated by pipetting 0.1 μL with a P2 micropipette under a Leica DMIL transmitted light microscope. Isolation was confirmed by pipetting the resulting droplet onto a microscopy slide and visually confirming the presence of single colonies, which were then transferred into 24-well plates containing 1 mL culture medium. Each well was seeded with 3 or 4 individual colonies. Wells were visually inspected 24 to 48 hours after transfer, and those containing colonies were transferred into a T25 culture flask containing 10 mL culture medium. Sheets were cultured in 1% to 10% Cereal Grass Medium (hereafter CGM3) in artificial seawater (hereafter ASW; as in (*84*)). Cultures were maintained at 22°C under a light-dark cycle (12:12 hours) in a VWR Scientific model 2005 low temperature incubator equipped with a lamp (Venoya Full Spectrum 150W Plant Growth LED) controlled by a programmable timer (Leviton VPT24-1PZ Vizia). For long-term storage, *C. flexa* cultures were frozen and kept in a liquid nitrogen dewar following (*85*). Cells could be successfully revived by thawing from frozen stocks (*85*).

#### Establishment and propagation of the ChoPs (*Choanoeca-Pseudomonas*) monoxenic strain

We define the ChoPs strain (Fig. 3B) as a monoxenic culture of *C. flexa* with the co-isolated bacterium *Pseudomonas oceani* (*P. oceani*). ChoPs was established by elimination of all environmental bacteria from the original polyxenic isolate by antiobiotic treatment, followed by re-seeding with *P. oceani*. More specifically, (1) a polyxenic isolate was treated with a combination of 6 antibiotics to prevent growth of environmental bacteria: carbenicillin (100 µg/mL), erythromycin (160 µg/mL), kanamycin (100 µg/mL), lincomycin (200 µg/mL), rifampicin (20 µg/mL), and streptamycin (200 µg/mL).

To confirm that the antibiotic combination prevented proliferation of all co-isolated bacteria, we inoculated 100 µL of the original polyxenic culture, with or without antibiotics, into 5 mL 1% CMG3 with 20 µg/mL cycloheximide (to prevent eukaryotic growth). Cells were grown in a bacterial incubator at 22°C under 300 rpm agitation and cell growth was monitored by measuring 600 nm optical density.

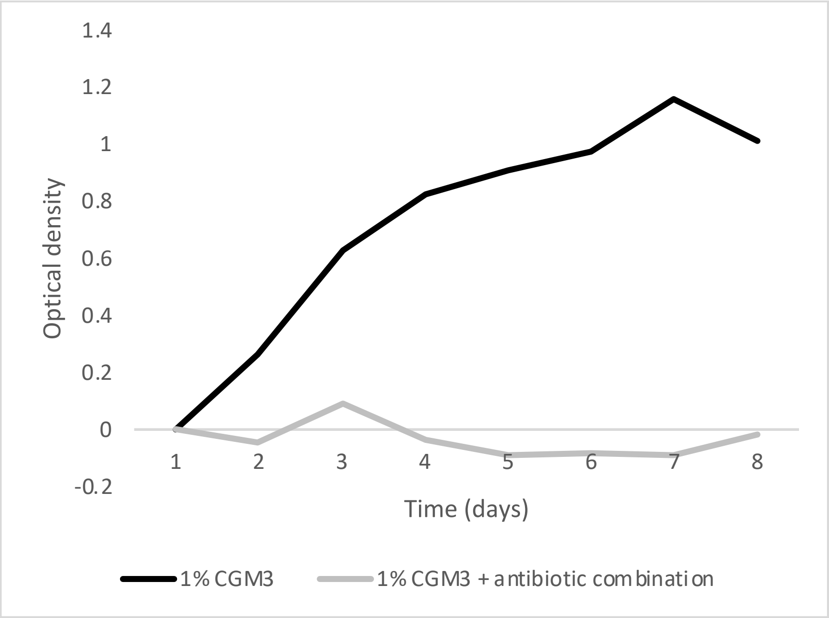


**Figure M1.** Growth of environmental bacteria co-isolated with *C. flexa*, monitored by optical density measurements, in the presence or in the absence of antibiotics.

To isolate *P. oceani*, 70 µL of the original polyxenic strain were spread on a 4% agar plate supplemented with CGM3 culture medium. A single colony was picked, grown in 5 mL 100% CGM3 for 3 days, and identified by 16S sequencing as *P. oceani*. We then passaged the antibiotic-treated *C. flexa* polyxenic line 5 times at 1:10 dilution in the 6-antibiotic combination, re-seeding every time with 1:20 volume of *P. oceani* culture (to progressively dilute out all other bacterial species, and feed choanoflagellates with a constant supply of pre-grown bacteria). After 5 passages, antibiotic treatment was replaced by treatment with 20 µg/mL rifampicin only (to which *P. oceani* is resistant) and active re-seeding with novel *P. oceani* was stopped. Monoxenicity of the resulting ChoPs line was validated by PCR amplification of the 16S locus followed by sequencing of 50 individual clones, as well as by iTag sequencing (Table S1 and section "Characterization of bacterial communities by iTag sequencing"). ChoPs was cultured in the constant presence of rifampicin to prevent contamination.

<u>*C. flexa* DNA extraction, rDNA locus amplification and phylogenetic analysis</u>

*C. flexa* was grown to $10^6$ cells/mL in 200 mL culture medium in 6-layer culture flasks, and pelleted by centrifugation for 15 minutes at 2,000 rpm at 4°C. DNA was extracted using the DNEasy Blood

& Tissue Kit (Qiagen 69504). The 18S locus was amplified by PCR with Q5 DNA Polymerase (New Englands Biolab M0491L) and the following degenerate primers:

| | **Forward primer** | **Reverse primer** |
|---|---|---|
| **First PCR** | CTCAARGAYTAAGCCATGCA | CCGCCCCAGYCAAACTCCC |
| **Nested PCR** | GAAACTGCGAATGGCTC | ACCTACGGAAACCTTGTTACG |

**Table M1.** Primer sequences for 18S cloning.

The following PCR programs were used: (1) for the first round of PCR: 1. 98 °C, 2 minutes; 2. 98 °C, 10 seconds; 3. 48 °C, 30 seconds; 4. 55 °C, 30 seconds; 5. 72 °C, 3 minutes; 6. repeat steps 2-5, 30x total; 7. 72 °C, 11 minutes 11 seconds; (2) for the nested PCR: 1. 98 °C, 2 minutes; 2. 98 °C, 10 seconds; 3. 48 °C, 30 seconds; 4. 52 °C, 30 seconds; 5. 72 °C, 1 minute; 6. repeat steps 2-5, 35x total; 7. 72 °C, 7 minutes 21 seconds.

rDNA sequences from *C. flexa* and other opisthokonts were aligned with ClustalX 2.0 (*86*), trimmed with gBlocks (http://phylogeny.lirmm.fr/phylo_cgi/one_task.cgi?task_type=gblocks) under minimally stringent parameters, and a Maximum Likelihood phylogenetic tree was produced with SeaView v. 4.7 (*87*) using PhyML with the GTR model, empirical nucleotide equilibrium frequencies, optimized proportion of invariable sites, optimal rate variation across sites, and best of NNI & SPR tree search method. Trees were visualized and edited with FigTree v. 1.4.4 (http://tree.bio.ed.ac.uk) and further edited with Adobe Illustrator CC 2018.

*C. flexa* RNA extraction, transcriptome sequencing and assembly

*C. flexa* was grown to $10^6$ cells/mL in 200 mL culture medium in 6-layer culture flasks and pelleted by centrifugation for 15 minutes at 2,000 rpm at 4°C. Cells were lysed by selective lysis as in (*71*) and RNA was extracted using a RNEasy kit (Qiagen 7404). 150 paired-end RNAseq libraries were prepared after poly-A selection by the QB3 Functional Genomics Laboratory at UC Berkeley with poly-A selection and sequenced on a HiSeq 4000 sequencer (Illumina, San Diego, California, United States) at the Vincent J. Coates Genomics Sequencing Laboratory at the California Institute for Quantitative Biosciences (Berkeley, California, United States). Read quality was assessed with FastQC (https://www.bioinformatics.babraham.ac.uk/projects/fastqc/) and transcriptome assembly was performed as in (*9*) with Trinity v. 2.5.1, with the '--trimmomatic' option and '-min_contig_length' set to 150. Predicted protein sequences were generated with Transdecoder (*88*) with a predicted minimum protein sequence length of 50 aminoacids and redundant protein sequences were eliminated using CD-HIT (*89*). The reads generated were uploaded onto the NCBI website with the BioSample accession number SAMN11533889, BioProject ID PRJNA540068, and SRA accession number PRJNA540068. Transcriptome and non-redundant predicted proteome are available on Figshare at DOI: 10.6084/m9.figshare.8216291. Phylogenetic trees for proteins of interest were generated following the same procedure as for rDNA (see above '*C. flexa* genome extraction, rDNA locus amplification and phylogenetic analysis') with aminoacid sequences instead of nucleotide sequences.

Characterization of bacterial communities by iTag sequencing

A *C. flexa* culture was pelleted at 4500xg and bacterial DNA was extracted using a DNeasy Kit (Qiagen, Hilden, Germany) following the manufacturer's Gram-Positive Bacteria Protocol. Library construction and sequencing were then performed by the UC Berkeley Functional Genomics Laboratory. Amplicon library construction was carried out in two distinct PCR steps: briefly, PCR1 used modified gene-specific primers to create amplification products of the V5-V6 region of the bacterial 16S gene flanked by stub sequences (to provide priming templates for PCR2); PCR2 primers added dual-matched index sequences, sequencing primer binding sites, and Illumina p5 and p7 adapter sequences to the 5' and 3' ends of each amplicon.
PCR1 was carried out in 25 µl reactions including 1 µl genomic DNA, 0.5 µl of each 10 µM primer, 10 µl 2.5x 5PRIME HotMasterMix (QuantaBio, Beverly, MA), 1µl BSA (New England Biolabs, Ipswitch, MA) (to a final concentration of 10µg/µL) and 12 µl nuclease free water. We amplified the 16S V4 hypervariable region using the primer set 515f from Parada et. al (*90*) (GTG**Y**CAGCMGCCGCGGTAA) and 806r from Apprill et al. (*91*) (GGACTAC**N**VGGGTWTCTAAT). All PCR reactions were set up on ice and using Hot Start polymerase master mix to minimize non-specific amplification and primer dimerization. PCR conditions were: denaturation at 94°C for 3 mins; 30 amplification cycles of 45 sec at 94°C, 1 min at 50°C and 90 sec at 72°C; followed by a 10 min final extension at 72°C. PCR products were visualized for successful amplification and correct sizing using gel electrophoresis.
PCR2 was carried out in 25 µl reactions including 5 µl genomic DNA, 2 µl of combined, µM forward and reverse indexing primers, 10 µl 2.5x 5PRIME HotMasterMix (QuantaBio, Beverly, MA), and 8 µl nuclease free water. PCR2 primers added dual--matched 8bp unique barcodes to each end of the amplicons, such that each forward and reverse primer pair carried the same index sequence. PCR conditions were: denaturation at 94°C for 3 mins; 8 amplification cycles of 45 sec at 94°C, 1 min at 52°C and 90 sec at 72°C; followed by a 10 min final extension at 72°C. PCR products were visualized and quantified using an Advanced Analytical Fragment Analyzer (Agilent, Santa Clara, CA), and individual samples were pooled equimolarly based on Fragment Analyzer concentrations.
The final pool of all individually indexed amplicon libraries was cleaned with Agencourt AMPure XP magnetic beads using a 0.8X bead ratio. The cleaned pool was quantified with qPCR using the KAPA Illumina Library Quant Kit and Universal qPCR Mix (KAPA Biosystems, Wilmington, MA). Amplicon libraries were sequenced on an Illumina MiSeq 300PE v3 run spiked with 10% PhiX in order to achieve sufficient sample heterogeneity.

Light microscopy

Sheets were imaged in FluoroDishes (World Precision Instruments FD35-100) by differential interference contrast (DIC) microscopy using a 40x (water immersion, C-Apochromat, 1.1 NA), 63x (oil immersion, Plan-Apochromat, 1.4 NA), or 100x (oil immersion, Plan-Apochromat, 1.4 NA) Zeiss objective mounted on a Zeiss Observer Z.1 with a Hamamatsu Orca Flash 4.0 V2 CMOS camera (C11440-22CU).

Transmission electron microscopy (TEM)

Sheets were concentrated by centrifugation (200xg for 5 min) and then resuspended in 5% BSA in artificial seawater. The resuspended sheets were then high pressure frozen using a Leica EM PACT2 and fixed by freeze substitution in 0.01% $OsO_4$ + 0.2% uranyl acetate in acetone. Samples were resin embedded in Epon Araldite (Embed-812) (*75*), cut into 80 nm sections, and then imaged using an FEI Tecnai 12 transmission electron microscope.

Scanning electron microscopy (SEM)

*C. flexa* sheets were concentrated by pelleting 6 mL of culture for 15 minutes at 200 g and gently resuspending the pellet in 600 µL final volume. Sheets were then pipetted onto silicon wafers coated with poly-D-lysine (Sigma Aldrich P6407-5MG), fixed for 2 hours in 2% glutaraldehyde in 0.1M Sodium cacodylate buffer pH 7.2, rinsed 3 times (for 15 minutes each) in 0.1M sodium cacodylate buffer pH 7.2 (for 15 minutes each), post-fixed for 2 hours in 1% osmium tetroxide in 0.1M sodium cacodylate buffer pH 7.2, and rinsed 3 times (for 5 minutes each) in 0.1M sodium cacodylate buffer, pH 7.2. Samples were dehydrated in the following steps: 35% ETOH (5 min), 50% ETOH (5 min), 70% ETOH (5 min), 80% ETOH (10 min), 95% ETOH (10 min), 100% ETOH (10 min) and 100% ETOH (10 min). Samples were critical point dried for 60 minutes on a Tousimis AutoSamdri 815 Critical Point Dryer, mounted on stubs using conductive carbon tape, and sputter coated before imaging on a Hitachi S-5000 Scanning electron microscope.

Sheet fixation and FM143-FX/phalloidin/immunofluorescence stainings

FluoroDishes (World Precision Instruments FD35-100) were pre-treated with a handheld Corona surface treater (Electro-Technic Products BD-20AC), coated with poly-D-lysine (Sigma Aldrich P6407-5MG) diluted to the provider's specifications, and rinced twice in ASW. Sheet colonies were then transferred into the treated dishes and left to adhere to the bottom surface for 30 minutes. Fixation was performed by adding 1:3 volume ice-cold 16% paraformaldehyde (PFA; reference) to the FluoroDish (to a final concentration of 4%) for 2 hours at room temperature.

**FM143-FX staining:** sheets were fixed with 5 µg/mL FM143-FX together with PFA in the fixation solution. This procedure was chosen because FM143-FX was observed to quickly trigger dissociation of live sheets, but to be compatible with the preservation of fixed samples. The fixation solution was then washed out three times carefully with ASW and the samples were imaged directly in FluoroDishes.

**Phalloidin staining:** cells were stained with 0.66 units/mL Alexa 488-phalloidin (ThermoFischer Scientific A12379) or rhodamine-phalloidin (ThermoFischer Scientific R415) in 0.3% Triton X/ASW overnight and under agitation at 4°C. The fixation solution was then washed out three times carefully with ASW and the samples were imaged directly in FluoroDishes.

**Immunofluorescence:** samples were blocked for 30 minutes at room temperature with blocking solution (1% bovine serum albumin in PEM buffer (100 mM PIPES pH 6.9, 1 mM EGTA, 1 mM MgSO4)/0.3% Triton-X), and then stained with antibodies diluted in blocking buffer (see below).

For antibodies against the heavy chain of myosin (or full-length myosin), we selected antibodies that were reported to bind both smooth and striated myosin isoforms, based on the following reasoning: as the divergence between both myosin paralogs predates the choanoflagellate/animal divergence, we expected any antibody that broadly targets both smooth and striated myosin of animals to be likely to bind to the choanoflagellate myosin heavy chain as well. Additionally, we investigated antibodies targeting the myosin regulatory light chain (either in all its forms or in its phosphorylated form only), motivated by its high level of sequence conservation (>80% identity between *C. flexa* and *H. sapiens*) and by conservation of the Ser19 phosphorylation residue. The following antibodies were tested:

| **Provider** | **Antibody** | **Target** |
|---|---|---|
| Developmental Hybridoma Bank | CMII23 | Myosin heavy chain |
| Sigma Aldrich | M7648 | Full-length myosin |
| Abgent | AP19667a-ev | Myosin regulatory light chain |
| Cell Signalling | 3671S | pSer19-Myosin Regulatory Light Chain |
| Bioss Inc. | bs-3295R | pSer19-Myosin Regulatory Light Chain |
| Sigma Aldrich | F1804-50UG | FLAG (negative control) |

Primary antibodies were diluted in blocking solution to the manufacturer's specifications. Primary antibody incubation was performed overnight at 4°C under agitation. Samples were then washed three times in PEM buffer, incubated with secondary antibodies diluted 1:300 in blocking solution (Goat anti-Mouse IgG1 Cross-Adsorbed Secondary Antibody, Alexa Fluor 488) for 2 hours at room temperature, washed three times with PEM, and imaged directly in FluoroDishes.

All samples were imaged using a Zeiss LSM 880 AxioExaminer with Airyscan and a 63x, 1.4 NA C Apo oil immersion objective (Zeiss) and excitation provided by a 488 nm laser (Zeiss).

Light-induced sheet inversion assays

Light-controlled inversion assays were performed in 24-well plates (Fischer Scientific 09-761-146) containing 2 mL sheet culture per well and imaged with a Leica DMIL LED transmitted light microscope coupled to a Leica DFC 350FX camera. Images were acquired with the MicroManager software at a frame rate of 1 image/second. The "light on" condition corresponded to maximal illumination by the microscope lamp and imaging with 1 ms exposure time. The histogram of detected light intensities was monitored with MicroManager and was found, under these conditions, to be a narrow bell curve centered around 50% light intensity. The "light off" condition was established by raising the exposure time to 400 ms and manually decreasing light intensity until the histogram of detected light intensities closely matched the one of the "light on" condition (sharp peak at 50%) – thus indicating that light intensity had been reduced down to close to 1:400 of its initial value. Each sheet was imaged for 120 seconds (55 to 60 seconds with the light off and 60 to 65 seconds after the light was switched off). The light reduction procedure itself lasted about 1 second (resulting in one white slice (see Movie S5) which was excluded from the area quantification pipeline – see below). This protocol was found to reliably induce inversion of polyxenic sheets during the sheets' subjective day.

Sheet area quantification and image analysis

Sheet projected area was quantified in ImageJ 1.46r. Stacks produced by live imaging of sheets undergoing light-induced inversion were cropped around the sheet of interest, split into 2 parts (before and after switching off the light, to correct for minimal differences in exposure), and processed with the "Make Binary" command followed by 2 to 5 iterations of the "Close" command and "Dilate" command in succession. Iterations were stopped when the resulting shape contained no gaps, and were performed an identical number of times for both parts of each movie. Areas were quantified by measuring the mean gray value on each slice with the measureStack plugin (http://www.optinav.info/MeasureStack.htm). Resulting measurements were stored in .xls files and combined into a single table using a Python or R script (*76*). Area=f(time) curves were plotted using the ggplot2 package in R studio (https://ggplot2.tidyverse.org/). Measurements were smoothened using a rolling average over a 5-second time window. For each individual sheet, areas were normalized by either (1) for the "light on" time window, the time average of areas measured over the entire window or (2) the initial area (for the "light off" time window).

Drug treatments and all-trans-retinal rescue assays

| **Name (target)** | **Stock concentration** | **Working concentration** |
|---|---|---|
| IBMX (PDE) | 1 M in DMSO | 1 mM |
| Caffeine (PDE) | 77 mM in $H_2O$ | 5 mM |
| Y-27632 (ROCK kinase) | 14 mg/mL in $H_2O$ | 43.7 µM |
| ML-7 (MRLC kinase) | 10 mg/mL in DMSO | 22 µM |
| All trans-retinal | 88 mM in ethanol | 88 µM |
| Latrunculin B (F-actin) | 20 mM in DMSO | 2-20 µM |
| Blebbistatin (myosin II) | 17 mM in 90%DMSO | 20 µM |
| 8-Br-cAMP | 100 mg/mL in $H_2O$ | 1E-5M to 1E-3M |
| 8-Br-cGMP | 50 mg/mL in $H_2O$ | 1E-5M to 1E-3M |

Sheets were pre-treated with actomyosin and ion channel inhibitors for 30 minutes before behavioral assays. Y-27632 did not observably affect inversion. EGTA was observed to instantly prevent sheet inversion after addition and, on long time scales (<30 minutes), led to sheet dissociation. All behavioral assays with EGTA were performed <1 minute after addition in the culture medium. Finally, caffeine, IBMX, and 8-Br-cGMP were observed to induce inversion in a few seconds after addition (if the culture medium was actively mixed by gentle swirling). For quantification of the number of inverted sheets, sheets were fixed by addition of 1:3 volume ice-cold 16% PFA, and manually counted under a Leica DMIL LED transmitted light microscope.

Bacterial rescue protocol

*Isolation of bacteria from monoxenic and polyxenic cultures*: To isolate bacteria from the monoxenic ChoPs culture *(P. oceani)* or polyxenic culture ("env. bacteria"), 20 µl of either culture was inoculated into 100% CGM3 with 20 µg/ml cycloheximide (to inhibit eukaryotic growth). Cultures were grown shaking at 22 degrees for 48 hours. Bacteria were then pelleted at 20,000xg, washed with ASW, and resuspended to 1 O.D.600/ml in ASW.

*Pre-treatment of recipient culture:* To reduce bacterial load, the recipient ChoPs culture was treated for 24 hours with an antibiotic cocktail (20 µg/ml rifampicin, 200 µg/ml streptomycin, 100 µg/ml kanamycin, 160 µg/ml erythromycin, 100 µg/ml carbenicillin, and 200 µg/ml lincomycin). After 24 hours, choanoflagellates were pelleted at 1000xg, washed once in ASW, then resuspended in 10%CGM3.
*Bacterial rescue:* Pre-treated Chops cultures were seeded into 10% CGM3 at 4E4 cells/ml; isolated bacteria (either *P. oceani* or "env. bacteria") were diluted into the culture at 1:1000 and grown for 48 hours. Cultures that received *P. oceani* were subsequently treated with 20ng/ul rifampicin for one week to ensure absence of contaminants. Photic response was then quantified as described below (see "Light-induced sheet inversions assays").

Retinal rescue protocol

Monoxenic ChoPs culture was grown in 10% CGM3 + 20 ng/µl rifampicin + 0 nM, 125 nM or 500 nM all-trans-retinal. After one week treatment, photic response was quantified as described above (see "Light-induced sheet inversion assays").

Sheet dissociation

A dense (1E6 cells/mL) sheet culture was reconcentrated 10x by centrifugation (2,000 rpm for 15 min at 4°C) and dissociated by vortexing for 30 seconds followed by filtration through a Tisch Scientific syringe-top 5 µm filter (SPEC18191). Dissociated cells were immediately transferred into an Ibidi 8-well plate imaging chamber (80826) coated with poly-D-lysine and washed twice with ASW (200 µL of cell suspension per chamber).

Quantification of actin ring diameter

Imaging of phalloidin labeled samples (both single cells and intact sheets) for actin ring quantification was performed as detailed in the "Sheet fixation and FM143-FX/phalloding/immunofluorescence stainings" section. Actin ring quantification for both intact and dissociated sheets was performed using Imaris v3.8 (Bitplane, Belfast). Images were first segmented with a global threshold chosen manually for each condition on one image and then held constant for all subsequent images. Diameters were measured using the "Measure Points" function, with three diameters manually chosen and the longest measured distance chosen as the ring diameter. Only completely intact rings with associated intact collars were chosen for analysis.

Sheet tracking

Sheets were imaged on a Zeiss Axio Zoom.V16 (generously lent by Zeiss to the 2018 Physiology course at the Marine Biology Laboratory in Woods Hole). Individual sheets were tracked on Fiji v. 2.0 (*82*) using the Manual Tracking option of the Tracking plugin.

Movement quantification

Movies of sheets swimming were produced on a Leica DMIL LED transmitted light microscope coupled to a Leica DFC 350FX camera, and movement was quantified by measuring inter-frame

Pearson correlation using RStudio (*83*). Sheet inversion was induced by a rapid (ca. 1 sec transition time) reduction in light intensity (see "Light-induced sheet inversion" section).

Microbead feeding assays, flow visualization and quantification

0.2 μm green fluorescent microbeads were used for both feeding assays and flow visualization (FluoSpheres™ Carboxylate-Modified Microspheres, 0.2 μm, red fluorescent (580/605), 2% solids ; ThermoFischer Scientific F8810) at a 1:100 dilution. For flow visualization, live sheets were observed in FluoroDishes on a Zeiss Z.1 Observer (see "Light microscopy" section). For feeding assays, 1 mL sheet culture were pipetted into wells in a 24-well plate with microbeads at 1:100 dilution, and put on a Reliable Inc. rocking shaker (see "Mechanically induced sheet inversion assays") under level 10 agitation for 1 hour. Sheets were then fixed by addition of 1:3 volume 16% PFA at 4°C (to a final concentration of 4% PFA) and carefully transferred into poly-D-lysine-coated FluoroDishes for observation, using a P1000 micropipette with a truncated pipet tip (to limit shear forces). Sheets were imaged on a Zeiss Z.1 Observer in both DIC (to observe cells) and green fluorescence (to reveal beads), and cells having phagocytosed beads were manually counted for each sheet.

Phototaxis assays and analysis

Phototaxis assays were performed in Ibidi 8-well plate imaging chambers (80826). Wells were completely filled with dense *C. flexa* cultures, and a coverslip (FisherScientific, 12-545-D) was used to cover the well in order to both prevent evaporation and flatten the background illumination intensity profile over the full extent of the well. Samples were then allowed to settle in ambient overhead light conditions for 30min. Next, a phase contrast reference image of the entire well was acquired using a Zeiss Z.1 Observer and Hamamatsu Orca Flash 4.0 V2 CMOS camera (C11440-22CU) by tiling with a 10x, 0.45 NA Plan-Apochromat (Zeiss) objective. The room was then darkened by turning off or blocking all sources of light. Except for the negative control condition, samples were then immediately exposed to directional illumination supplied by a white LED (Elegoo) powered by an Elegoo Uno R3 Arduino supplying 5V through across a 10 kΩ (Elegoo) resistor and focused onto one edge of the well containing the sample using a 15x magnification lens (TV-15 Triview, Carson Optical). This illumination scheme created a conical gradient of decreasing light intensity along the well in the direction of illumination due to increasing distance from the focal point at the edge of the well. Negative controls received no directional illumination. After one hour, a final image of the entire well was acquired as before. For single cell phototaxis experiments, prior to the settling step, sheets were dissociated by vortexing in a 1.5 mL Eppendorf tube using a Vortex Genie 2 (Scientific industries) for one minute.

All image processing was performed using Fiji (*82*). First, images were background subtracted using Gaussian blurred ($\sigma$ = 100 pixels) duplicate images. Images were then cropped to remove the edges of well, and a five-pixel median filter was applied. Images were then imported into MATLAB release 2016a (Mathworks, Natick) where subsequent analysis was performed. For each pair of reference and final images (see "Phototaxis assays" section), a phototaxis index (PI) was calculated, with PI defined as:

$$PI = \frac{I_{pf}}{I_{df}} - \frac{I_{pr}}{I_{dr}}$$

where $I_{pr}$ is the integrated intensity of the first 1/3 of the reference image proximal to the phototaxis light source, $I_{dr}$ is the integrated intensity of the 1/3 of the reference image most distal to the phototaxis light source, $I_{pf}$ is the integrated intensity of the first 1/3 of the final image proximal to the phototaxis light source, and $I_{df}$ is the integrated intensity of the 1/3 of the final image most distal to the phototaxis light source. Because choanoflagellate cells are bright under phase contrast, PI is a readout of the relative change in abundance of choanoflagellate cells in the region proximal versus distal to the light source over the course of the phototaxis assay.

Live imaging of *Diaphanoeca grandis*, *Salpingoeca rosetta*, *Salpingoeca urcoelata* and *Monosiga brevicollis*

*D. grandis*, *S. rosetta*, *S. urceolata* and *M. brevicollis* cultures were obtained by thawing frozen stocks stored in liquid nitrogen in the King lab (following (*85*)) and maintained in 1% CGM3/ASW medium. Live cells were mounted in FluoroDishes treated with a handheld Corona surface treater, coated with poly-D-lysine, and rinsed three times with ASW. Imaging was performed in DIC optics at 100x magnification on a Zeiss Observer Z.1 platform using a Hamamatsu Orca Flash 4.0 V2 CMOS camera (C11440-22CU).

**Supplementary Figures**

**Fig. S1. Full 18S rDNA phylogenetic tree of *C. flexa,* several other choanoflagellates, and representative outgroup opisthokonts (including animals and fungi).**

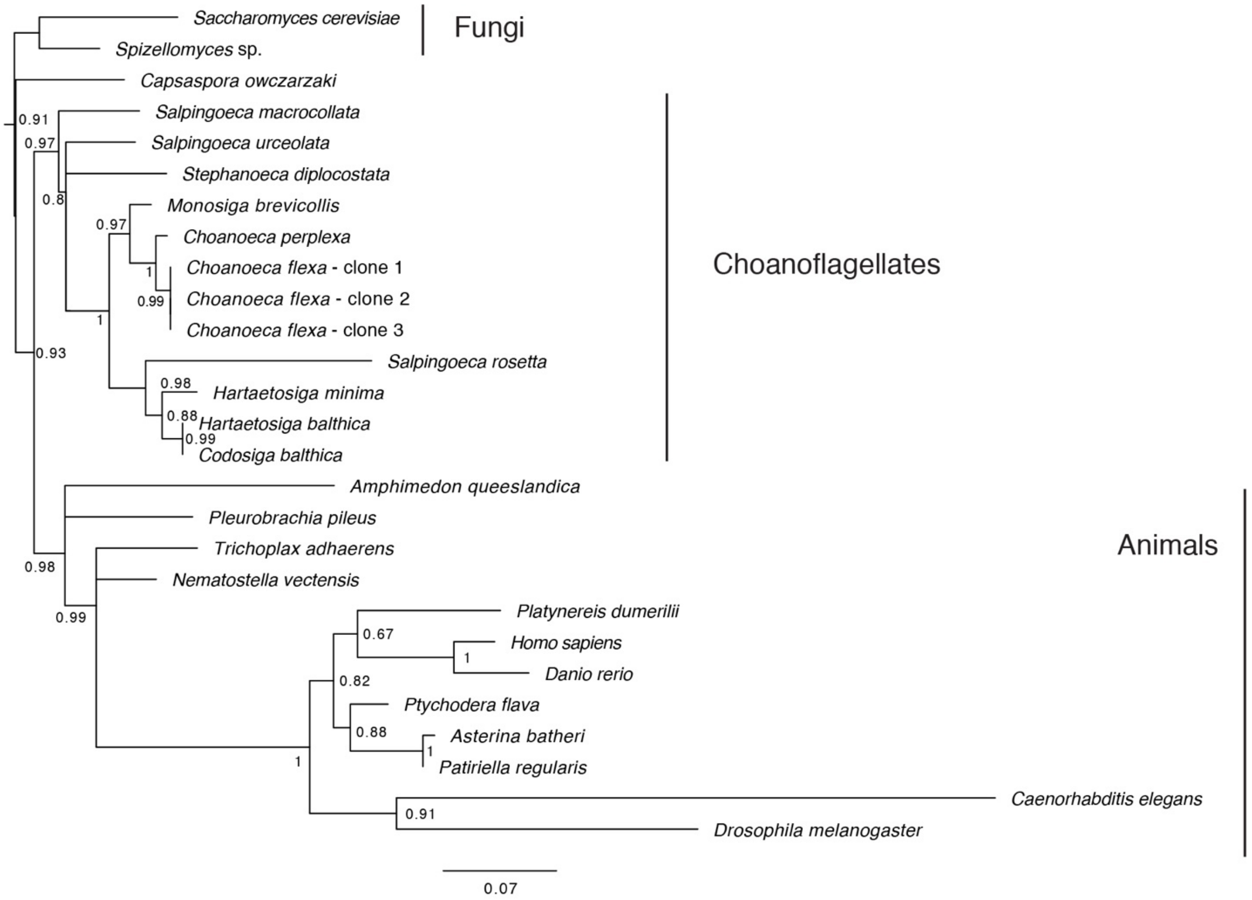


Maximum Likelihood phylogenetic tree of 18S rDNA sequences from *C. flexa* and other opisthokonts. Support values on nodes: approximate likelihood ratio test (aLRT) (*77*). Nodes with support values <0.5 were collapsed into polytomies (e.g. the branching order between the sponge *Amphimedon queeslandica*, the ctenophore *Pleurobrachia pileus* and other metazoans, which represents a currently uncertain point of higher-order metazoan phylogeny (*78*)).

**Fig. S2. *C. flexa* encodes four putative homologs of RhoPDE.**

**A**

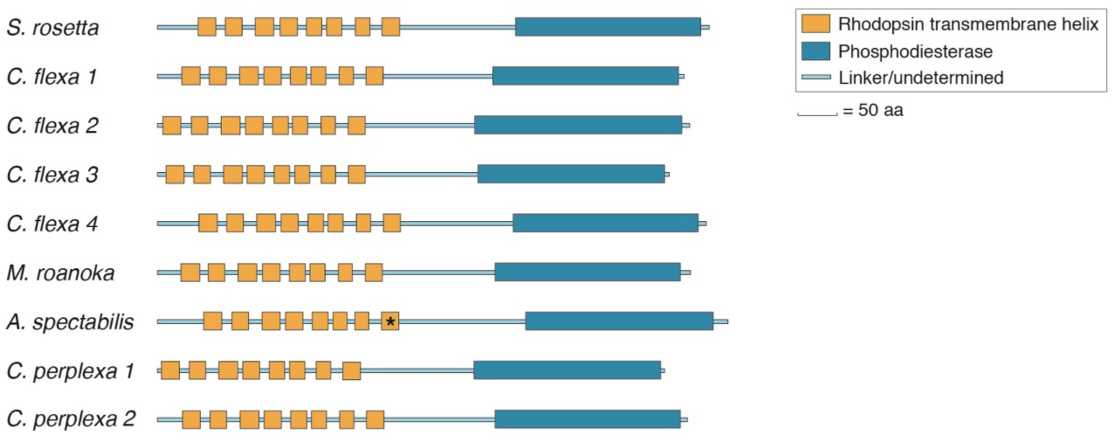


**B**

| | | | | | | | | | | | | | | |
|---|---|---|---|---|---|---|---|---|---|---|---|---|---|---|
| | **No. in *S. rosetta* RhoPDE** | 161 | 164 | 165 | 168 | 172 | 175 | 264 | 267 | 268 | 271 | 292 | 295 | 296 |
| | No. in BR | 82 | 85 | 86 | 89 | 93 | 96 | 182 | 185 | 186 | 189 | 212 | 215 | 216 |
| | **Transmembrane Helix** | 4 | 4 | 4 | 4 | 4 | 4 | 7 | 7 | 7 | 7 | 8 | 8 | 8 |
| | ***S. rosetta* RhoPDE** | R | E | W | T | L | W | W | F | P | E | D | A | K |
| | ***C. flexa* RhoPDE 1** | R | E | W | T | L | W | W | F | P | E | D | A | K |
| | ***C. flexa* RhoPDE 2** | R | E | W | T | L | W | W | H | P | E | D | A | K |
| | ***C. flexa* RhoPDE 3** | R | E | W | T | L | V | W | F | P | E | D | A | K |
| | ***C. flexa* RhoPDE 4** | R | E | W | T | L | W | W | F | P | E | D | A | K |
| H+ pump | BR | R | D | W | T | L | D | W | Y | P | W | D | A | K |
| Cl- pump | HR | R | T | W | S | I | A | W | Y | P | W | D | A | K |
| Na+ pump | KR2 | R | N | W | D | L | Q | W | Y | P | Y | D | S | K |
| cation channel | ChR2 | R | E | W | T | I | H | W | F | P | F | D | S | K |
| anion channel | ACR1 | R | S | W | T | M | L | W | Y | P | W | D | C | K |
| sensor | SRII | R | D | W | T | I | F | W | Y | P | W | D | T | K |
| | ASR | R | D | W | T | L | S | W | Y | P | W | P | S | K |
| guanylate cyclase | Rhod-GC | R | E | W | T | L | L | W | F | P | W | D | A | K |
| | Rhod-GC Allomyces | R | E | R | T | L | V | W | V | P | A | D | A | K |
| histidine kinase | HKR1 | R | M | W | T | M | L | W | F | P | W | D | G | K |
| | HKR (cop6) | R | Q | W | T | M | I | W | F | P | W | N | A | K |

**C**

| | | | | | | | | | | | | | | |
|---|---|---|---|---|---|---|---|---|---|---|---|---|---|---|
| | **No. in *S. rosetta* RhoPDE** | 492 | 605 | 609 | 616 | 620 | 623 | 624 | 644 | 656 | 657 | 660 | 661 | 693 |
| | No. in PDE10A | 564 | 674 | 678 | 685 | 689 | 692 | 693 | 713 | 725 | 726 | 729 | 730 | 762 |
| | ***S. rosetta* RhoPDE** | D | D | P | N | S | L | Q | F | G | E | F | I | W |
| | ***C. flexa* RhoPDE 1** | D | D | P | T | A | L | Q | F | G | E | F | I | W |
| | ***C. flexa* RhoPDE 2** | D | D | P | A | A | L | Q | F | G | E | F | I | Y |
| | ***C. flexa* RhoPDE 3** | N | D | N | A | A | L | Q | F | A | Q | H | F | W |
| | ***C. flexa* RhoPDE 4** | D | D | P | T | A | L | Q | F | G | E | F | I | W |
| | 1B | D | D | P | H | T | L | M | L | S | Q | F | I | W |
| | 2A | D | D | Q | T | A | I | Y | M | L | Q | F | M | W |
| Both | 3A | D | D | P | H | T | I | V | F | L | Q | F | I | W |
| | 10A | D | D | V | T | A | I | Y | M | G | Q | F | Y | W |
| | 11A | D | D | V | S | A | V | T | I | L | Q | W | I | W |
| | 4A | D | D | P | Y | T | I | M | M | S | Q | F | I | A |
| cAMP | 7A | D | D | P | S | S | V | T | L | I | Q | F | M | W |
| | 8B | D | D | P | C | A | I | S | V | S | Q | F | I | W |
| | 5A | D | D | I | Q | A | V | A | L | M | Q | F | I | W |
| cGMP | 6A | D | D | I | Q | A | V | A | M | L | Q | F | I | W |
| | 9A | D | D | E | A | V | L | L | F | A | Q | F | I | Y |

**(A)** Domain architectures of RhoPDE proteins across choanoflagellates. Shown are all choanoflagellate genes found to contain a phosphodiesterase domain and the 7 helices characteristic of bacterial (type I) rhodopsins. Phosphodiesterase domains were annotated using PFAM. Rhodopsin transmembrane helices of the S. rosetta protein were annotated following Lamarche et al. (*22*); those of the other homologs were determined by alignment with the S. rosetta protein. *The 8th transmembrane helix of A. spectabilis RhoPDE lacks the conserved lysine residue required to covalently bind retinal, and therefore likely lacks rhodopsin activity. **(B** and **C)** Comparison of conserved amino acids between *S. rosetta* RhoPDE, *C. flexa* RhoPDEs, and various type I rhodopsins and phosphodiesterases. Proteins and residues were selected as described by Yoshida *et al.* (*21*). Red and blue letters indicate acidic and basic amino acids, respectively. **(B)** Comparison of conserved amino acids between choanoflagellate RhoPDEs and selected type I rhodopsins. Notably, all of the *C. flexa* homologs contain the conserved lysine residue of the 8th transmembrane helix, which is required for covalent binding of the chromophore retinal. **(C)** Comparison of conserved amino acids between choanoflagellate RhoPDEs and selected cyclic nucleotide phosphodiesterases. The substrate specificities of the phosphodiesterases are indicated in the leftmost column. *S. rosetta* RhoPDE hydrolyzes cGMP ~10-fold more efficiently than cAMP based on *in vitro* studies (*21*).

**Fig. S3. Neither *C. flexa* nor the bacterium *Pseudomonas oceani* encodes the complete retinal biosynthesis pathway.**

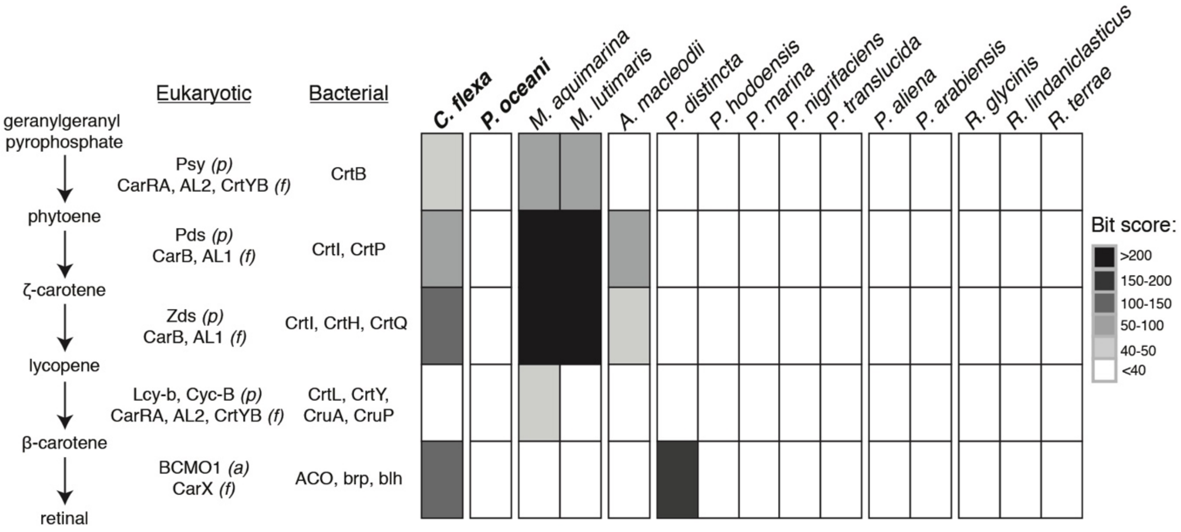


The *C. flexa* transcriptome, as well as the genomes of the bacterial species found in the polyxenic culture (Table S1), were searched for genes encoding enzymes in the retinal biosynthesis pathway using BLASTP. For each step in the pathway, multiple bacterial, plant ("p"), fungal ("f"), and/or animal ("a") genes were used as queries, and the highest returned score is shown. Notably, *C. flexa* does not encode a recognizable lycopene cyclase, the enzyme that synthesizes beta-carotene, the immediate precursor of retinal. Thus, *C. flexa* must receive retinal or beta-carotene from its food. *Pseudomonas oceani,* the bacterium in the monoxenic ChoPs culture, lacks any recognizable homologs of genes in this pathway, indicating that it likely cannot produce beta-carotene or retinal. The *Bordetella* sp. bacterium shown in Table S1 was not included in this analysis because the precise species could not be identified.

**Fig. S4. Phylogenetic trees of predicted proteins encoded by the *Choanoeca flexa* transcriptome.**

A

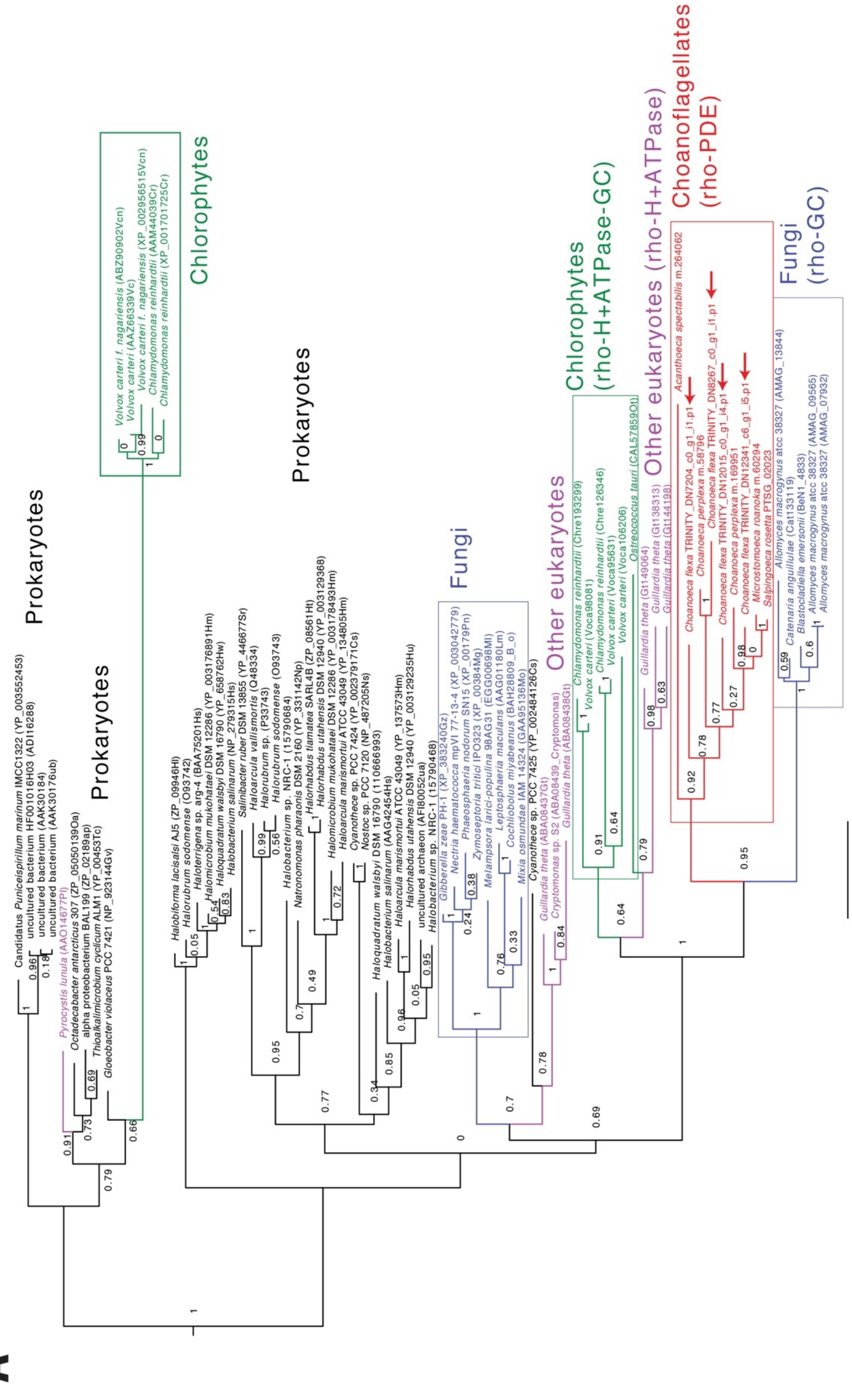

B

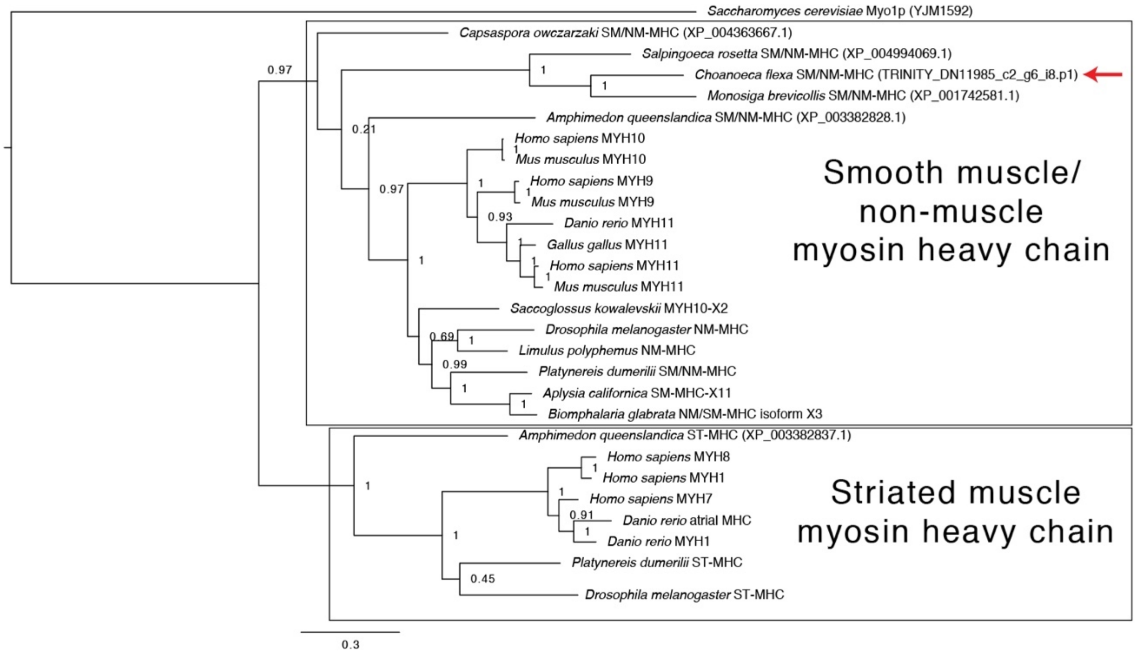
Saccharomyces cerevisiae Myo1p (YJM1592)
Capsaspora owczarzaki SM/NM-MHC (XP_004363667.1)
Salpingoeca rosetta SM/NM-MHC (XP_004994069.1)
Choanoeca flexa SM/NM-MHC (TRINITY_DN11985_c2_g6_i8.p1)
Monosiga brevicollis SM/NM-MHC (XP_001742581.1)
Amphimedon queenslandica SM/NM-MHC (XP_003382828.1)
Homo sapiens MYH10
Mus musculus MYH10
Homo sapiens MYH9
Mus musculus MYH9
Danio rerio MYH11
Gallus gallus MYH11
Homo sapiens MYH11
Mus musculus MYH11
Saccoglossus kowalevskii MYH10-X2
Drosophila melanogaster NM-MHC
Limulus polyphemus NM-MHC
Platynereis dumerilii SM/NM-MHC
Aplysia californica SM-MHC-X11
Biomphalaria glabrata NM/SM-MHC isoform X3
Amphimedon queenslandica ST-MHC (XP_003382837.1)
Homo sapiens MYH8
Homo sapiens MYH1
Homo sapiens MYH7
Danio rerio atrial MHC
Danio rerio MYH1
Platynereis dumerilii ST-MHC
Drosophila melanogaster ST-MHC
Smooth muscle/
non-muscle
myosin heavy chain
Striated muscle
myosin heavy chain
0.97
0.21
0.97
0.93
0.69
0.99
0.91
0.45
0.3

C

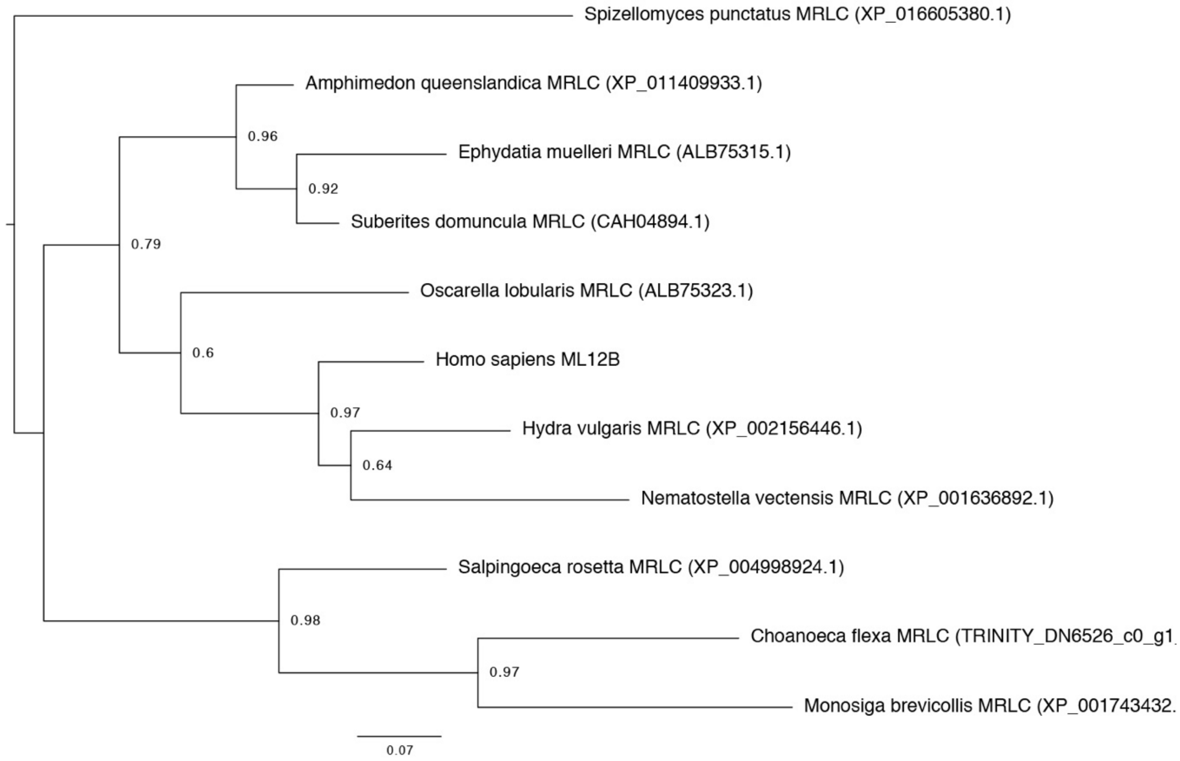
Spizellomyces punctatus MRLC (XP_016605380.1)
Amphimedon queenslandica MRLC (XP_011409933.1)
Ephydatia muelleri MRLC (ALB75315.1)
Suberites domuncula MRLC (CAH04894.1)
Oscarella lobularis MRLC (ALB75323.1)
Homo sapiens ML12B
Hydra vulgaris MRLC (XP_002156446.1)
Nematostella vectensis MRLC (XP_001636892.1)
Salpingoeca rosetta MRLC (XP_004998924.1)
Choanoeca flexa MRLC (TRINITY_DN6526_c0_g1
Monosiga brevicollis MRLC (XP_001743432.
0.96
0.92
0.79
0.6
0.97
0.64
0.98
0.97
0.07

**(A)** Phylogenetic tree of microbial rhodopsin domains, based on sequences compiled by Avelar et al. (*70*) to which sequences from five choanoflagellates with microbial rhodopsins (*C. flexa* (red arrows) and sequences from Richter et al. (*9*) – see Fig. S2) were added. Choanoflagellate microbial rhodopsins are all fused to phosphodiesterase domain (see Fig. S4) and are sister to a clade of fungal microbial rhodospins fused to a guanylyl-cyclase, in agreement with the phylogenetic tree reconstructed by Avelar et al (*70*). **(B)** Phylogenetic tree of myosin heavy chains (MHC). The *C. flexa* transcriptome appears to encode a single predicted MHC homolog (red arrow) which, like in other studied choanoflagellates (*40*, *55*), belongs to the smooth muscle/non-muscle family. **(C)** Phylogenetic tree of myosin regulatory light chains (MRLC). The *C. flexa* transcriptome appears to encode a single predicted MRLC homolog.

**Fig. S5. Sheets with flagella out are more motile than sheets with flagella in.**

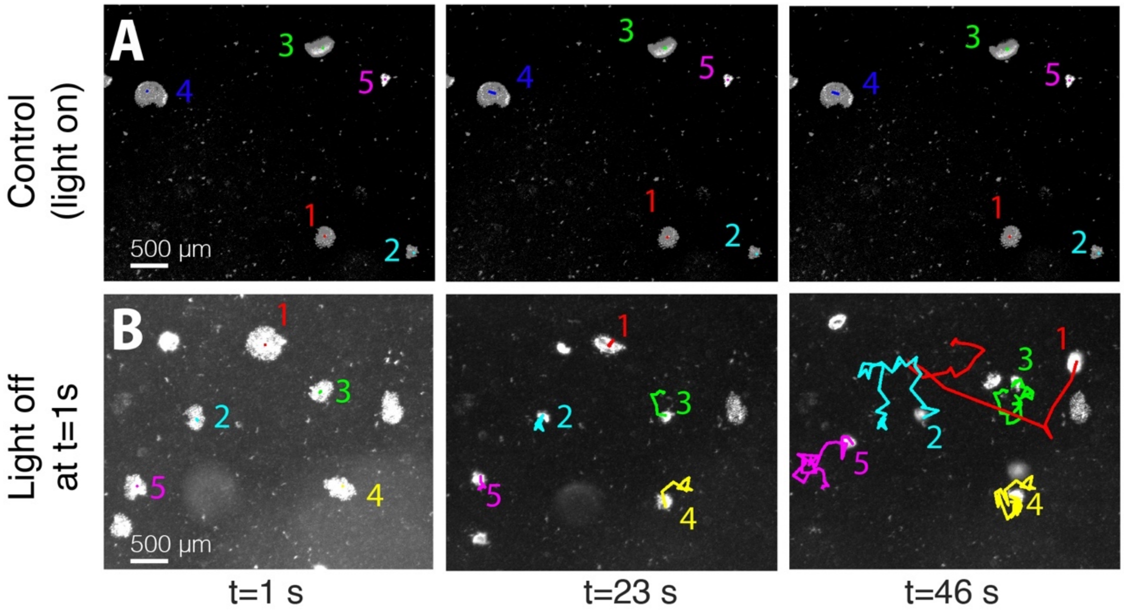


**(A)** Five sheets observed at low magnification in a constant level of transmitted light (frames from Movie S6). Sheets remained flagella-in and nearly immotile. Colored curves represent tracks of individual sheets. **(B)** Five different sheets observed in transmitted light following a rapid (~1 sec) reduction in light intensity (frames from Movie S7). Following inversion, the sheets became actively motile. Colored lines represent tracks of individual sheets.

**Fig. S6. The flow field generated by flagellar beating is reoriented during sheet inversion.**

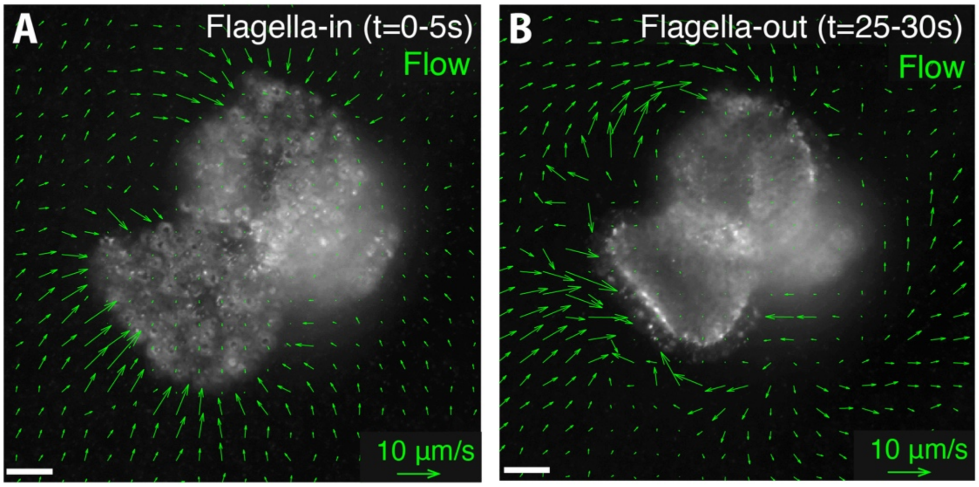


**(A)** Feeding flow generated by a sheet with flagella in (visualized by observation of 0.2 µm fluorescent beads in the water and computed by Particle Image Velocimetry; see Material and Methods). The flow is predominantly directed toward the center of the sheet, consistent with it carrying bacterial prey toward the cell bodies. **(B)** Flow generated by the same sheet after inversion. Note that the flow is predominantly directed parallel to border of the sheet, making it unlikely to support efficient feeding.

**Fig. S7. Sheet phototaxis requires retinal and multicellularity.**

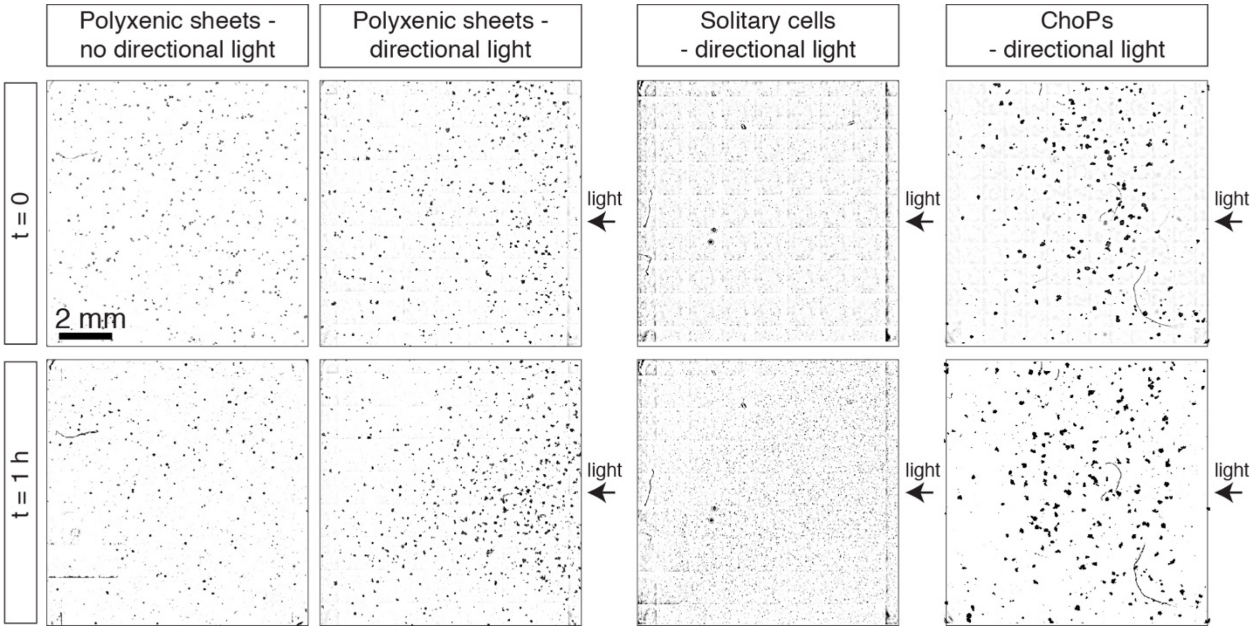


Top row: sheets imaged at low magnification at t=0 by transmitted light. Bottom row: sheets imaged at t=1 hour by transmitted light. Polyxenic sheets (cultured with a diversity of co-isolated bacterial species) under directional illumination accumulated toward the side of the flask near the light source (on the right side of the image), while no directional accumulation was observed in ChoPs monoxenic sheets (without retinal and unable to perceive light), polyxenic sheets without directional illumination, and single cells.

**Fig. S8. Neighboring cells within sheets are linked by their collars.**

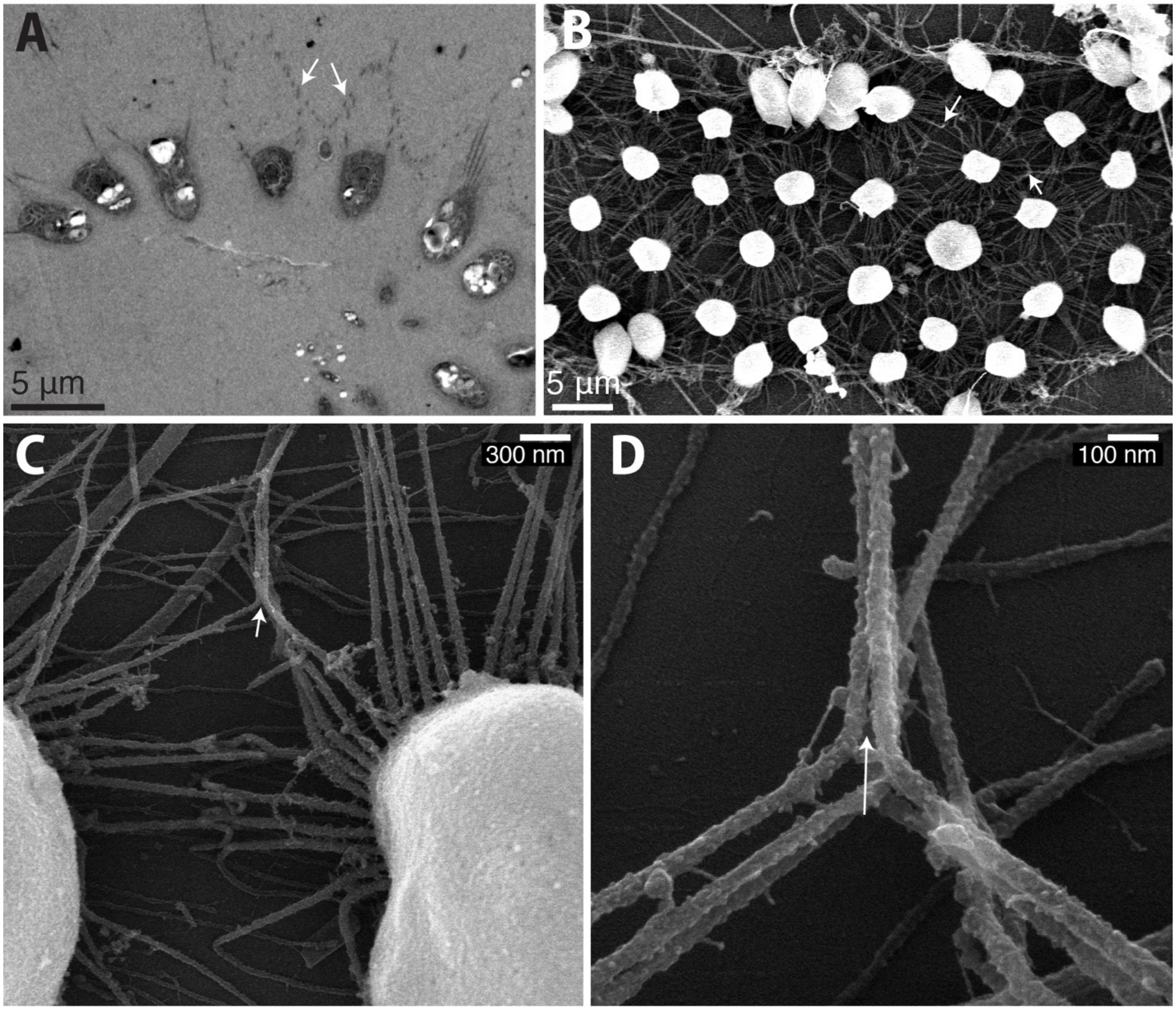


**(A)** Cross-section of a sheet imaged by transmitted electron microscopy. Note the close proximity between neighboring collars (arrows), and the absence of visible basal extracellular matrix, basal filopodia, or intercellular bridges. **(B)** Sheet imaged by scanning electron microscopy (SEM). Collars are spread out on the EM disk with cell bodies sticking up. Note the direct contact between neighboring collars (arrows) and the lack of any other structure connecting neighboring cells. **(C)** Contact between microvilli (arrow) belonging to two neighboring cells imaged by SEM. **(D)** Same individual as (**C**) imaged at higher magnification. Note the close apposition of the microvilli belonging to two neighboring cells (arrow), which appear closer to each other than microvilli are within the collar of the same cell.

**Fig. S9. Caffeine treatment results in cell-autonomous collar deformations.**

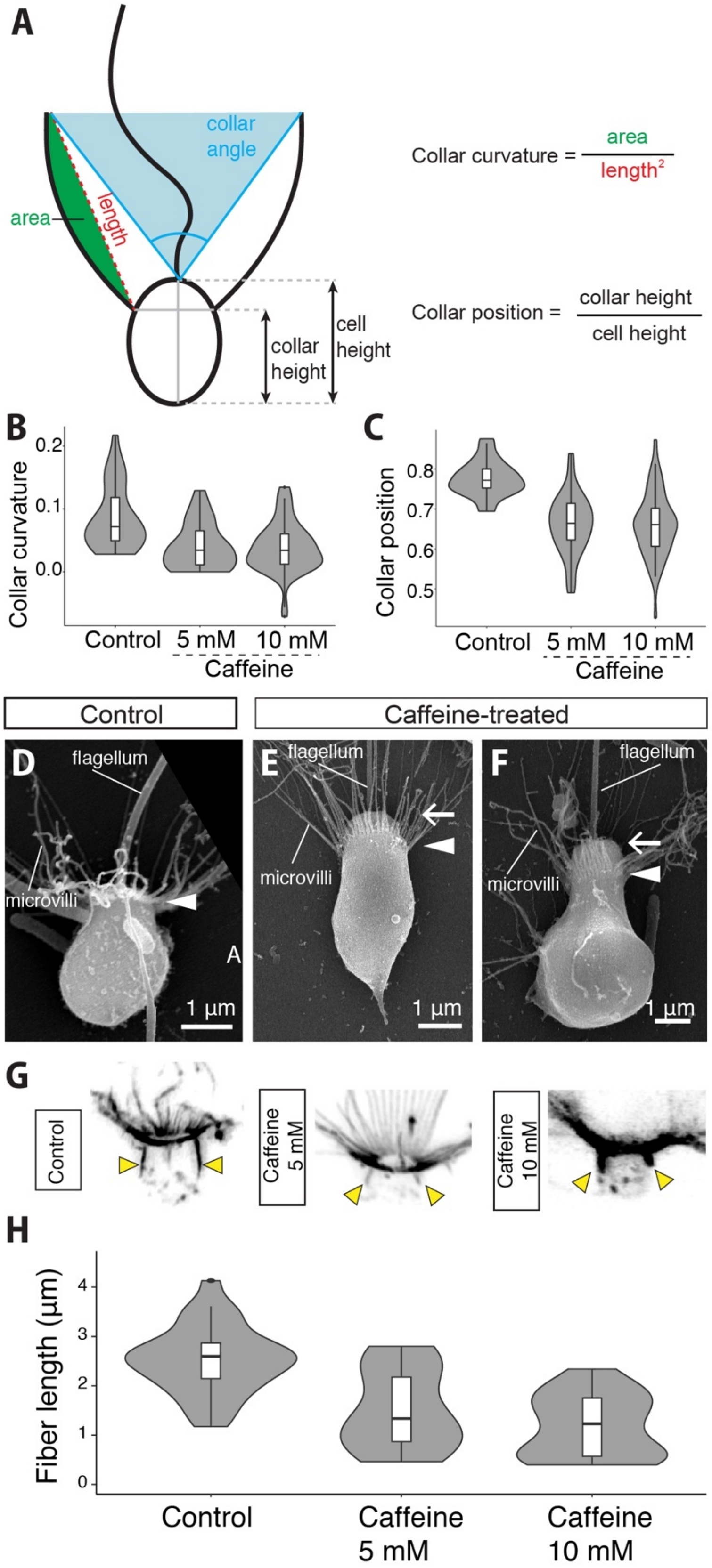

**(A)** Schematic summarizing the values measured and calculated to quantify collar shape. **(B)** Caffeine treatment leads to a decrease in collar curvature in dissociated cells. $n = 30$ untreated cells, $n = 35$ cells treated with 5 mM caffeine, and $n = 59$ cells treated with 10 mM-caffeine. $p = 4.6*10^{-5}$ (control vs. 5 mM caffeine) and $p = 2.9*10^{-6}$ (control vs. 10 mM caffeine) by Dunnett's test for comparing several treatments with a control. **(C)** Caffeine treatment leads to sliding of the collar toward the basal pole in dissociated cells. $n = 27$ untreated cells, $n = 35$ cells treated with 5 mM caffeine, and $n = 57$ cells treated with 10 mM caffeine. $p = 1.6*10^{-8}$ (control vs. 5 mM caffeine) and $p = 5.7*10^{-5}$ (control vs. 10 mM caffeine) by Dunnett's test. **(D** to **F)** Untreated (**D**) and caffeine-treated (**E** and **F**) dissociated cells. In untreated cells, the base of the collar (arrowhead) appears close to the apical side of the cell (arrow; where the flagellum emerges), while it is significantly displaced toward the base in caffeine-treated cells. Note the "bottle cell"-like morphology of the cell in (**F**) (with a narrow apex and a bulbous base), which is characteristic of a fraction of caffeine-treated cells (or cells in intact sheets during inversion) and is considered diagnostic of apical constriction in animal cells (*43*, *79*, *80*). **(G** and **H)** Longitudinal actin (yellow arrowheads) fibers shorten in response to caffeine treatment in dissociated cells. **(G)** Cells stained for F-actin by fluorescent phalloidin. Yellow arrowheads: longitudinal fibers. **(H)** Fiber length quantification. $n = 28$ control cells, $n = 16$ cells treated with 5 mM caffeine, and $n = 20$ cells treated with 10 mM caffeine. $p = 5.2*10^{-5}$ (control vs. 5 mM caffeine) and $p = 3.9*10^{-8}$ (control vs. 10 mM caffeine) by Dunnett's test.

**Fig. S10. Five distinct anti-myosin II antibodies stain the apical actin ring in *Choanoeca flexa*.**

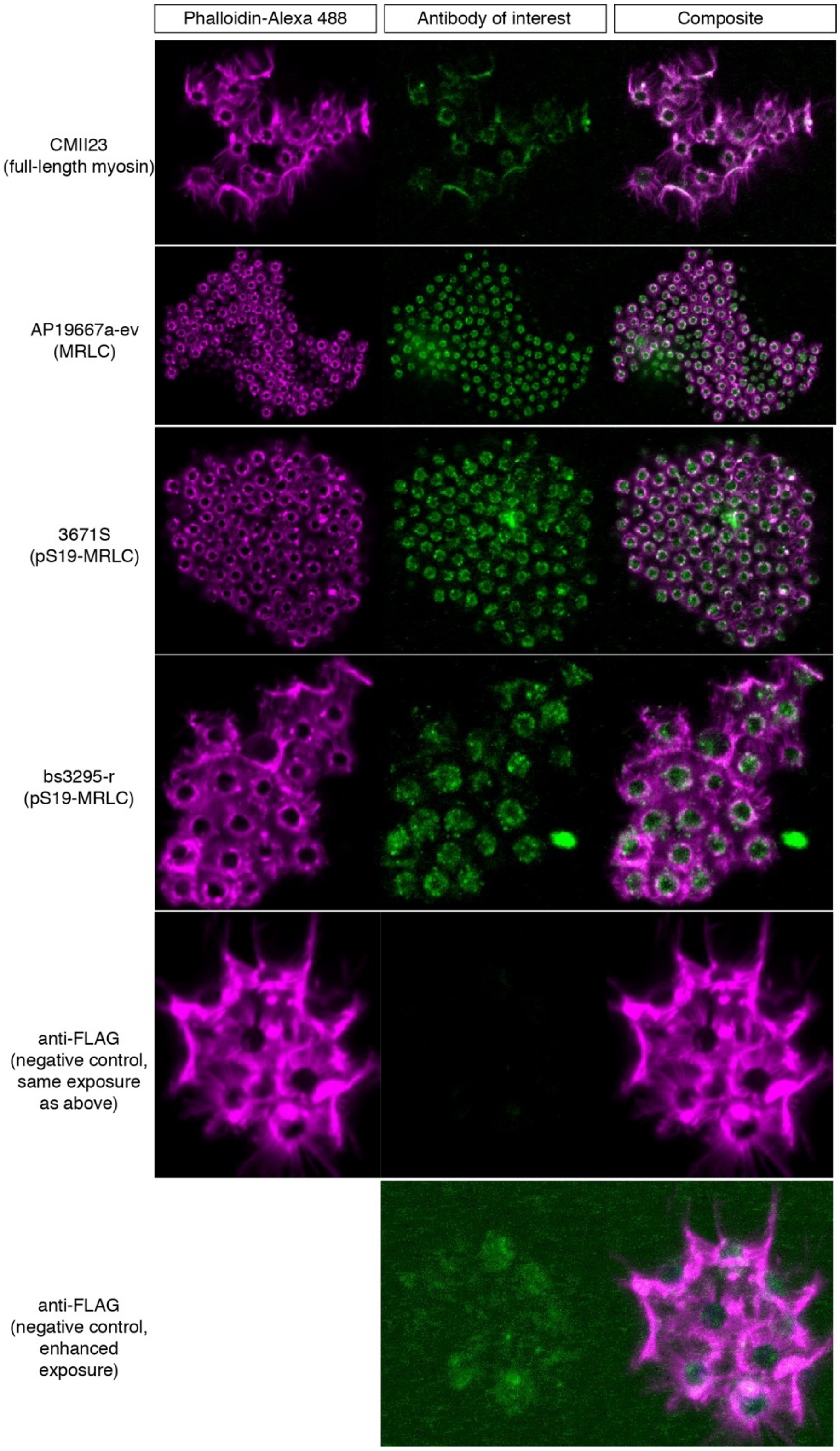

Sheets immunostained by the antibody of interest (Developmental Hybridoma Bank CMII23 against full-length myosin; Abgent AP19667a-ev against Myosin Regulatory Light Chain; and Cell Signalling Technology 3671S and Bioss USA bs-3295r against phosphorylated pS19-Myosin Regulatory Light Chain) and co-stained with rhodamine-phalloidin (targeting F-actin). All antibodies stain a domain nested in (and partly overlapping with) the apical actin ring, while a negative control antibody (Sigma Aldrich F1804 anti-FLAG) gives no detectable pattern under the same imaging conditions and rendering parameters. In sheets stained with the negative control antibody, only unspecific staining in the cell body is apparent if one increases the brightness of the corresponding channel with ImageJ.

**Fig. S11. Myosin II is required for apical ring constriction in response to caffeine.**

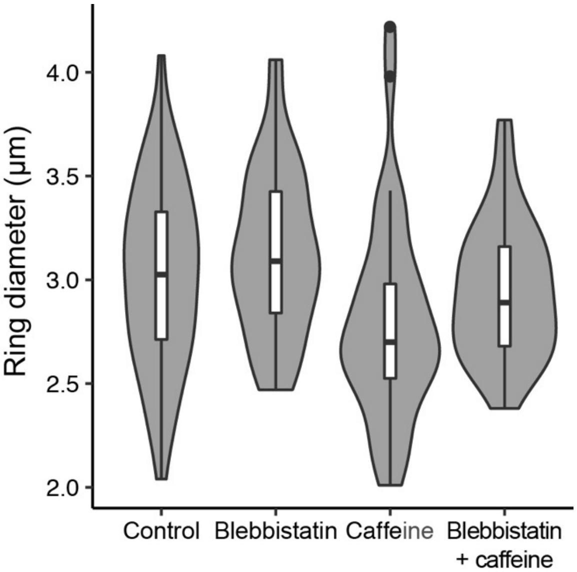


Ring diameter measured in dissociated cells, untreated, treated with 2 µM blebbistatin (which inhibits myosin II), treated with 5 mM caffeine, or treated with both 2 µM blebbistatin and 5 mM caffeine. Blebbistatin treatment prevents caffeine-induced ring constriction. $n$ = 164 control cells, $n$ = 106 blebbistatin-treated cells, $n$ = 77 caffeine-treated cells, and $n$ = 99 cells treated with both blebbistatin and caffeine, respectively. $p$ = 0.0111 (caffeine vs. control), $p$ = 0.0034 (caffeine vs. blebbistatin) and $p$ = 0.0097 (caffeine vs. blebbistatin+caffeine) by Kruskal-Wallis test for multiple comparisons. Other comparisons show no significant difference: $p$ = 1.00 (blebbistatin vs. control and blebbistatin vs. blebbistatin+caffeine) and $p$ = 0.76 (blebbistatin vs. blebbistatin+caffeine).

**Fig. S12. Spontaneous deformations of the collar in the choanoflagellates *Diaphanoeca grandis* and *Salpingoeca rosetta*.**

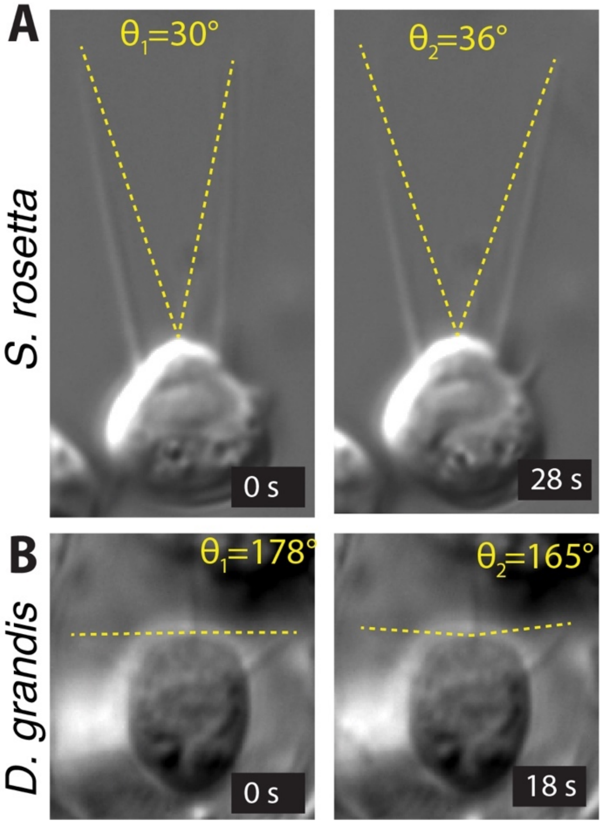


Time-lapse DIC imaging of live *S. rosetta* cells shows spontaneous reorientation of individual microvilli (Movie S10), notably following an increase in light intensity. In the loricate *D. grandis*, which is encased in a self-secreted silicon-based extracellular lodge, spontaneous and reversible changes in microvilli curvature are observed (Movie S11). Both types of collar deformation can be quantified as a change in collar angle (yellow doted lines).

**Figure S13. Phototransduction pathways in choanoflagellates, animals, and fungi.**

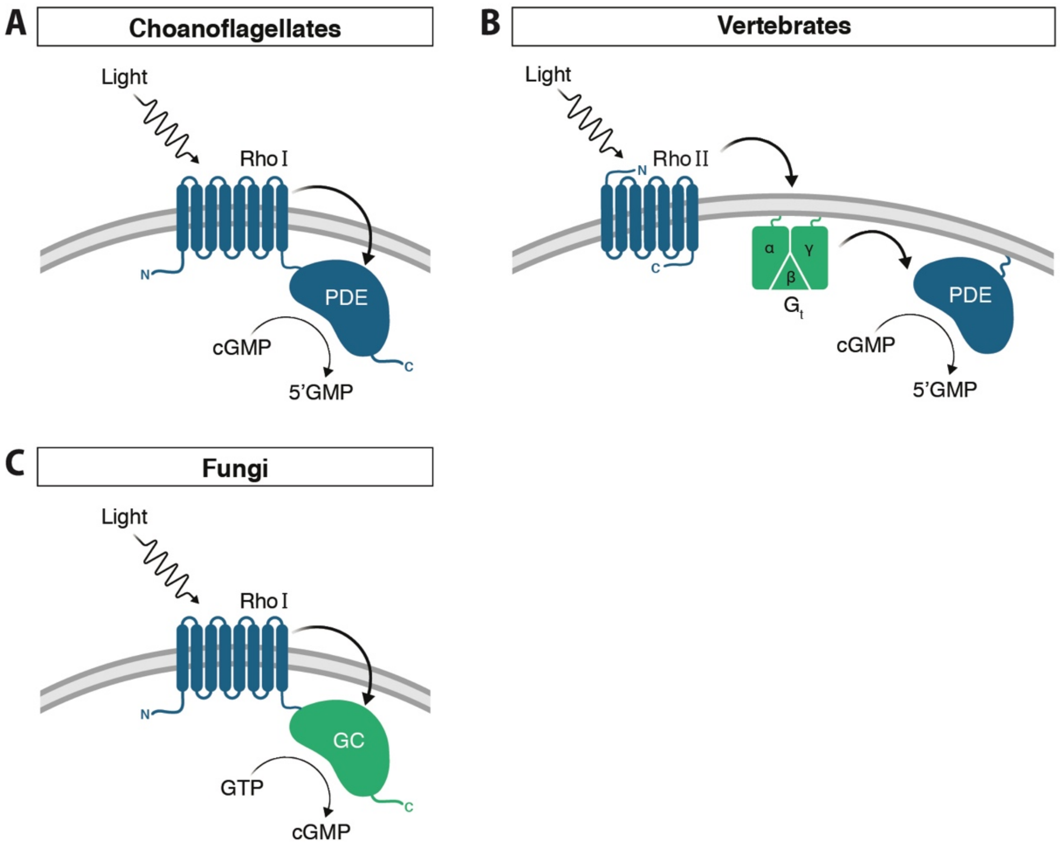


**(A)** Many choanoflagellates (Fig. S2) encode a fusion protein, RhoPDE, composed of a type I (bacterial) rhodopsin fused to a cyclic nucleotide phosphodiesterase (PDE). When illuminated, the rhodopsin activates the PDE domain, resulting in hydrolysis of cGMP (*20*, *21*).
**(B)** A similar rhodopsin-cGMP pathway controls phototransduction in vertebrate photoreceptor cells. Light activates a type II (eukaryotic) rhodopsin in the disk membrane, which activates a cGMP-specific phosphodiesterase via a G-protein intermediary (Gt). The G-protein and the PDE are tethered to the membrane by lipid modifications (*29*, *63*).
**(C)** Like choanoflagellates, some fungal zoospores use a type I rhodopsin fusion protein for phototransduction. However, this fungal protein, RhoGC, comprises a rhodopsin fused not to a PDE but to a guanylyl cyclase (GC), which catalyzes light-dependent synthesis of cGMP from GTP. Thus, illumination results in increased cellular cGMP levels (*70*).

**Supplementary Table**

**Table S1. Bacterial composition of polyxenic sheet culture determined by 16S sequencing.**

| Phylum | Species | Reads | Representative 16S Sequence |
|---|---|---|---|
| Gammaproteobacteria | *Pseudoalteromonas* sp.<br>*(distincta, hodoensis, marina, nigrifaciens,* or *translucida)* | 147,406<br>(57.1%) | AGCGTTAATCGGAATTACTGGGCGTAAAGCGTACGCAGGCGGTT<br>TGTTAAGCGAGATGTGAAAGCCCCGGGCTCAACCTGGGAACTGC<br>ATTTCGAACTGGCAAACTAGAGTGTGATAGAGGGTGGTAGAATT<br>TCAGGTGTAGCGGTGAAATGCGTAGAGATCTGAAGGAATACCGA<br>TGGCGAAGGCAGCCACCTGGGTCAACACTGACGCTCATGTACGA<br>AAGCGTG |
| Gammaproteobacteria | *Alteromonas macleodii* | 62,765<br>(24.3%) | AGCGTTAATCGGAATTACTGGGCGTAAAGCGCACGCAGGCGGTT<br>TGTTAAGCTAGATGTGAAAGCCCCGGGCTCAACCTGGGATGGTC<br>ATTTAGAACTGGCAGACTAGAGTCTTGGAGAGGGGAGTGGAATT<br>CCAGGTGTAGCGGTGAAATGCGTAGATATCTGGAGGAACATCAG<br>TGGCGAAGGCGACTCCCTGGCCAAAGACTGACGCTCATGTGCGA<br>AAGTGTG |
| Gammaproteobacteria | *Pseudomonas oceani* | 21,464<br>(8.3%) | AGCGTTAATCGGAATTACTGGGCGTAAAGCGCGCGTAGGCGGCT<br>TGATAAGATGGGTGTGAAATCCCCGGGCTTAACCTGGGAACTGC<br>ATCCATAACTGTCTGGCTAGAGTACAGTAGAGGGTGGTGGAATT<br>TCCTGTGTAGCGGTGAAATGCGTAGATATAGGAAGGAACACCAG<br>TGGCGAAGGCGACCACCTGGACTGATACTGACGCTGAGGTGCGA<br>AAGCGTG |
| Gammaproteobacteria | *Pseudoalteromonas aliena* | 1,6271<br>(6.3%) | AGCGTTAATCGGAATTACTGGGCGTAAAGCGTACGCAGGCGGTT<br>TGTTAAGCGAGATGTGAAAGCCCCGGGCTCAACCTGGGAACTGC<br>ATTTCGAACTGGCAAGCTAGAGTGTGATAGAGGGTGGTAGAATT<br>TCAGGTGTAGCGGTGAAATGCGTAGAGATCTGAAGGAATACCGA<br>TGGCGAAGGCAGCCACCTGGGTCAACACTGACGCTCATGTACGA<br>AAGCGTG |
| Bacteroidetes | *Muricauda* sp.<br>*(aquimarina* or *lutimaris)* | 5,419 (2.1%) | AGCGTTATCCGGAATCATTGGGTTTAAAGGGTCCGTAGGCGGGC<br>CTGTAAGTCAGGGGTGAAAGTTTGTGGCTCAACCATAAAATTGC<br>CTTTGATACTGCAGGTCTTGAGTCATGGTGGGGTTGCCGGAACA<br>TGTGGTGTAGCGGTGAAATGCATAGATATCACATAGAACACCGA<br>TCGCGAAGGCAGGTGACCAACCATGTACTGACGCTGATGGACGA<br>AAGCGTG |
| Gammaproteobacteria | *Pseudoalteromonas arabiensis* | 4,567 (1.8%) | AGCGTTAATCGGAATTACTGGGCGTAAAGCGTACGCAGGCGGTT<br>TGTTAAGCGAGATGTGAAAGCCCTGGGCTCAACCTGGGAACTGC<br>ATTTCGAACTGGCAAACTAGAGTGTGATAGAGGGTGGTAGAATT<br>TCAGGTGTAGCGGTGAAATGCGTAGAGATCTGAAGGAATACCGA<br>TGGCGAAGGCAGCCACCTGGGTCAACACTGACGCTCATGTACGA<br>AAGCGTG |
| Gammaproteobacteria | *Bordetella* sp. | 443<br>(0.171%) | AGCGTTAATCGGAATTACTGGGCGTAAAGCGTGCGCAGGCGGTT<br>CGGAAAGAAAGATGTGAAATCCCAGGGCTTAACCTTGGAACTGC<br>ATTTTTAACTGTCGAACTAGAGTGCGTCAGAGGGGGGTGGAATT<br>CCGCGTGTAGCAGTGAAATGCGTAGAGATGCGGAGGAACACCGA<br>TGGCGAAGGCAGCCCCCTGGGATGACACTGACGCTCATGCACGA<br>AAGCGTG |
| Gammaproteobacteria | *Rhodanobacter* sp.<br>*(lindaniclasticus, glycinis,* or *terrae)* | 10 (0.004%) | AGCGTTAATCGGAATTACTGGGCGTAAAGGGTGCGTAGGCGGTT<br>AGTTAAGTCTGTTGTGAAATCCCCGGGCTCAACCTGGGAATGGC<br>AATGGATACTGGCTAGCTAGAGTGTGTCAGAGGATGGTGGAATT<br>CCCGGTGTAGCGGTGAAATGCGTAGAGATCGGGAGGAACATCAG<br>TGGCGAAGGCGGCCATCTGGGACAACACTGACGCTGAAGCACGA<br>AAGCGTG |

The bacterial species present in the polyxenic *C. flexa* culture were identified by iTag sequencing of 16S rDNA (*81*). Shown are all species for which at least 10 reads were recovered. Due to the short length of the 16S amplicon (rightmost column), some reads were compatible with multiple bacterial species (e.g., first row). The 16S sequence of the *Bordetella* bacterium did not match any species in the NCBI database.

**Supplementary Movie Captions**

**Movie S1.** Sheet inversion and relaxation observed in an environmental splash pool sample.

**Movie S2.** Sheet inversion (from flagella-in to flagella-out) observed in DIC transmitted light microscopy.

**Movie S3.** Sheet relaxation (from flagella-out to flagella-in) observed in DIC transmitted light microscopy.

**Movie S4. Spontaneous sheet inversion, showing decrease in projected area.** Time lapse movie of a sheet observed in DIC transmitted light microscopy.

**Movie S5. Light-to-dark transitions reliably induce sheet inversion, which can be quantified as a decrease in projected area.** Left: time lapse movie of a sheet observed in phase contrast microscopy. Right: projected area. The light-to-dark transition is visible as a white frame at t=52 seconds, and corresponds to a 75-fold decrease in the intensity of incident light. Camera exposure time was increased correspondingly to allow continued imaging.

**Movie S6.** Sheets in constant light display have low mobility. Sheet culture observed with a Zeiss AxioZoom transmitted light dissecting microscope.

**Movie S7.** Light-to-dark transitions induces sheet inversion followed by fast swimming. Sheet culture observed with a Zeiss AxioZoom transmitted light dissecting microscope.

**Movie S8.** Spontaneous collar contractions in *Salpingoeca urceolata*. Cells are attached to the substrate by a stalked, cup-shaped extracellular lodge called a theca. The most spectacular contraction event is visible shortly after t=610 seconds and corresponds to a rapid decrease in collar angle, concomitant with retraction of the cell body inside the theca. The cell then reverts to its resting shape in the following 20 seconds.

**Movie S9.** Spontaneous collar contractions in *Monosiga brevicollis*. Bacteria co-cultured with *M. brevicollis* are visible around the cell body.

**Movie S10.** Spontaneous reorientation of microvilli in *Salpingoeca rosetta*.

**Movie S11.** Spontaneous changes in collar curvature in *Diaphanoeca grandis*. The cell is encased within an extracellular lodge called a lorica, composed of self-secreted silicon strips.

**Movie S12.** Spontaneous collar contractions in the thecate form of *Choaneca flexa*. Note that the thecate form lacks a flagellum, as in *C. perplexa* (*19*), the sister-species of *C. flexa*.
